# FUS controls muscle differentiation and structure through LLPS mediated recruitment of MEF2 and ETV5

**DOI:** 10.1101/2024.09.18.613669

**Authors:** Gina Picchiarelli, Anne Wienand, Salim Megat, Amr Aly, Marije Been, Nibha Mishra, Saskia Hutten, Erin Sternburg, Pierre Cauchy, Stéphane Dieterle, Marica Catinozzi, Valérie Demais, Laura Tzeplaeff, Annemarie Huebers, Dagmar Zeuschner, Angela Rosenbohm, Albert C. Ludolph, Anne-Laurence Boutillier, Tobias Boeckers, Dorothee Dormann, Maria Demestre, Chantal Sellier, Clotilde Lagier-Tourenne, Erik Storkebaum, Luc Dupuis

**Affiliations:** Université de Strasbourg, Inserm, Strasbourg Translational Neuroscience and Psychiatry, UMR-S1329, Centre de Recherches en Biomédecine; Strasbourg, France; Molecular Neurobiology Laboratory, Donders Center for Neuroscience, Donders Institute for Brain, Cognition and Behaviour and Faculty of Science, Radboud University; Nijmegen, Netherlands; Institute of Anatomy and Cell Biology, Ulm University; Ulm, Germany; Department of Neurology, The Sean M. Healey and AMG Center for ALS, Massachusetts General Hospital, Harvard Medical School, Boston, MA, USA; Institute of Molecular Physiology, Johannes Gutenberg-Universität (JGU), Mainz, Germany; Plateforme Imagerie In Vitro, CNRS UPS-3156, NeuroPôle, Strasbourg, France; Université de Strasbourg, UMR 7364 CNRS, Laboratoire de Neurosciences Cognitives et Adaptatives (LNCA), Strasbourg, France; Department of Neurology, Heinrich Heine-University, Düsseldorf, Germany; Electron Microscopy Unit, Max Planck Institute for Molecular Biomedicine, Münster, Germany; Department of Neurology, Ulm University; Ulm, Germany; German Center for Neurodegenerative Diseases (DZNE) Ulm; Ulm, Germany; Institute of Molecular Biology (IMB) Mainz, Germany

## Abstract

FUS is an RNA binding protein mutated in amyotrophic lateral sclerosis (ALS), a neurodegenerative disease characterized by progressive muscle weakness. We show that ALS-associated *FUS* mutations lead to ultrastructural defects in muscle of *FUS-*ALS patients, with disruption of sarcomeres and mitochondria. Studies in mouse and *Drosophila* models demonstrate an evolutionary-conserved cell autonomous function of FUS in muscle development. Mechanistically, FUS is required for transcription of MEF2 dependent genes, binds to the promoter of genes bound by ETS transcription factors in particular ETV5 and co-activates transcription of MEF2 dependent genes with ETV5. FUS phase separates with ETV5 and MEF2A, and MEF2A binding to FUS is potentiated by ETV5. Last, *Etv5* haploinsufficiency exacerbates muscle weakness in a mouse model of *FUS-*ALS. These findings establish FUS as an essential protein for skeletal muscle structure through its phase separation-dependent recruitment of ETV5 and MEF2, defining a novel pathway compromised in *FUS-*ALS.

## Introduction

Amyotrophic lateral sclerosis (ALS) is the most frequent adult-onset motor neuron disease, leading to death within a few years after onset of motor symptoms ^1–3^. ALS is a genetically heterogeneous disease, with currently more than 30 associated genes. Four genes (*C9ORF72, SOD1, FUS, TARDBP*) are the major causes of familial forms of ALS. Motor symptoms of ALS are associated with degeneration of motor neurons and progressive muscle atrophy and weakness. In *SOD1*-ALS, analyses of transgenic mouse models suggested that muscle weakness is primarily a consequence of motor neuron degeneration ^4^. Indeed, reducing mutant SOD1 expression in skeletal muscle of mutant SOD1 mice did not affect the disease course ^5,6^ and stimulating muscle mitochondrial biogenesis did not modify disease progression or survival ^7^ in spite of delayed muscle atrophy. On the other hand, restricted expression of mutant SOD1 in skeletal muscle led to skeletal muscle damage and altered motor neurons or neuromuscular junctions ^8–10^, suggesting a possible contribution of mutant SOD1 expression in skeletal muscle to ALS-related weakness.

Compelling evidence support the contribution of skeletal muscle defects in the pathogenesis of spinal and bulbar muscular atrophy (SBMA, a.k.a. Kennedy’s disease), another motor neurodegenerative disease caused by a polyglutamine repeat expansion in the androgen receptor (AR) ^11^. In a knock-in SBMA mouse model, myopathic and neurogenic skeletal muscle pathology preceded motor neuron pathology ^12^, and both male SBMA patients and female heterozygous carriers displayed myogenic and neurogenic myopathy ^13^. In addition, muscle-specific excision of the AR121Q transgene or its peripheral knockdown in a SBMA mouse model prevented motor phenotypes, muscle pathology and motor neuronopathy, and dramatically extended survival ^14,15^. Thus, expression of ALS-mutant genes in skeletal muscle might contribute to motor neuron disease to various extent, depending on the specific ALS gene ^4,16^ and the cell autonomous impact on muscles of ALS-associated genes other than SOD1 remains to be determined.

One of the most aggressive forms of ALS is caused by mutations in the *FUS* gene, encoding an RNA binding protein with prominent roles in transcription, splicing and local translation ^17,18^. *FUS-*ALS patients develop motor symptoms on average 10 to 20 years earlier than other ALS patients and show fast disease progression ^19–22^. *FUS* mutations cluster in the C-terminus of the FUS protein, encoding the nuclear localization signal, which results in impaired nuclear import of the FUS protein ^23^. The most severe phenotypes associated with *FUS* mutations are observed in patients carrying a frameshift mutation leading to C-terminal truncation of the protein and pronounced mislocalization of FUS to the cytoplasm ^19–21^. We and others previously generated knock-in mouse models for *FUS*-ALS, expressing a truncated FUS protein ^24–27^. *Fus*^ΔNLS/+^ mice show cytoplasmic accumulation of FUS in the central nervous system, similar to *FUS-*ALS patients ^24,25,28^. These mice develop mild muscle weakness, with late onset motor neuron degeneration and cognitive phenotypes ^24,25,28–30^. *FUS* is a ubiquitously expressed gene, including in skeletal muscle, and recent work showed abnormalities in energy metabolism pathways in *FUS*-ALS skeletal muscle ^31,32^, yet it remains unknown whether these defects are cell autonomously caused by expression of mutant FUS in skeletal muscle. Using mouse models and iPSC-derived myotubes of ALS patients, we recently showed that FUS is enriched in subsynaptic nuclei in skeletal muscles, and controls transcription of acetylcholine receptor subunit genes ^30^. This suggested that expression of mutant FUS in skeletal muscle may directly contribute to neuromuscular demise in this severe form of ALS.

Following up on these observations, we show here that FUS is required for muscle differentiation. Loss or mutation of *Fus* in muscle cells led to pronounced transcriptional defects, with a converging signature positioning MEF2 as a key disrupted transcription factor. Mechanistically, FUS directly binds to promoters of genes containing an ETV5-binding motif in muscle cells, and we demonstrate that FUS, MEF2A and ETV5 cooperate to activate transcription of their target genes. As a consequence, muscle ultrastructure, including mitochondrial ultrastructure, was severely compromised in *Fus^ΔNLS^* mice. Importantly, selectively reverting the *Fus^ΔNLS^* mutation to wild type in mouse skeletal muscle rescued ultrastructural defects, while muscle-selective loss of either FUS or ETV5 orthologs in Drosophila was sufficient to lead to muscle degeneration. Last, iPSC-derived myotubes and muscle biopsies of *FUS*-ALS patients showed similar prominent sarcomeric and mitochondrial ultrastructural defects, suggesting that disruption of FUS-dependent transcriptional pathways contribute to muscle defects in *FUS*-ALS patients.

## Results

### Ultrastructural and mitochondrial muscle defects in *FUS-*ALS patients and iPSC derived myotubes

To characterize the impact of *FUS* mutations on skeletal muscles, we first analyzed muscle biopsies from two healthy control individuals and 4 *FUS-*ALS patients (**Supplementary Table 1**). FUS protein was mostly nuclear in controls but mislocalized to the cytosol in *FUS-*ALS muscles (**Figure 1a**). Overall decrease of cytochrome c with accumulation at the plasma membrane was observed in *FUS-*ALS muscles by immunofluorescence staining (**Figure 1a**). Cytochrome c oxidase (COX) detected by histochemistry was also decreased in *FUS-*ALS muscles compared to controls (**Figure 1b**). We then performed transmission electron microscopy (TEM) on the same muscle biopsies. While healthy control muscle showed normal structure and properly aligned Z-lines, muscles from all four *FUS-*ALS patients showed myofibrillar disorganization and Z-line abnormalities. Quantification demonstrated a high percentage of fragmented, curvy and out of alignment *zigzag* Z-lines in all *FUS-*ALS muscles (**Figure 1c**). We also observed significant defects in mitochondrial ultrastructure in *FUS-*ALS muscles, including fragmentation, swelling or complete destruction of mitochondria in 3 out of the 4 ALS*-FUS* patients (**Supplementary** Figure 1a).

**Figure 1:**
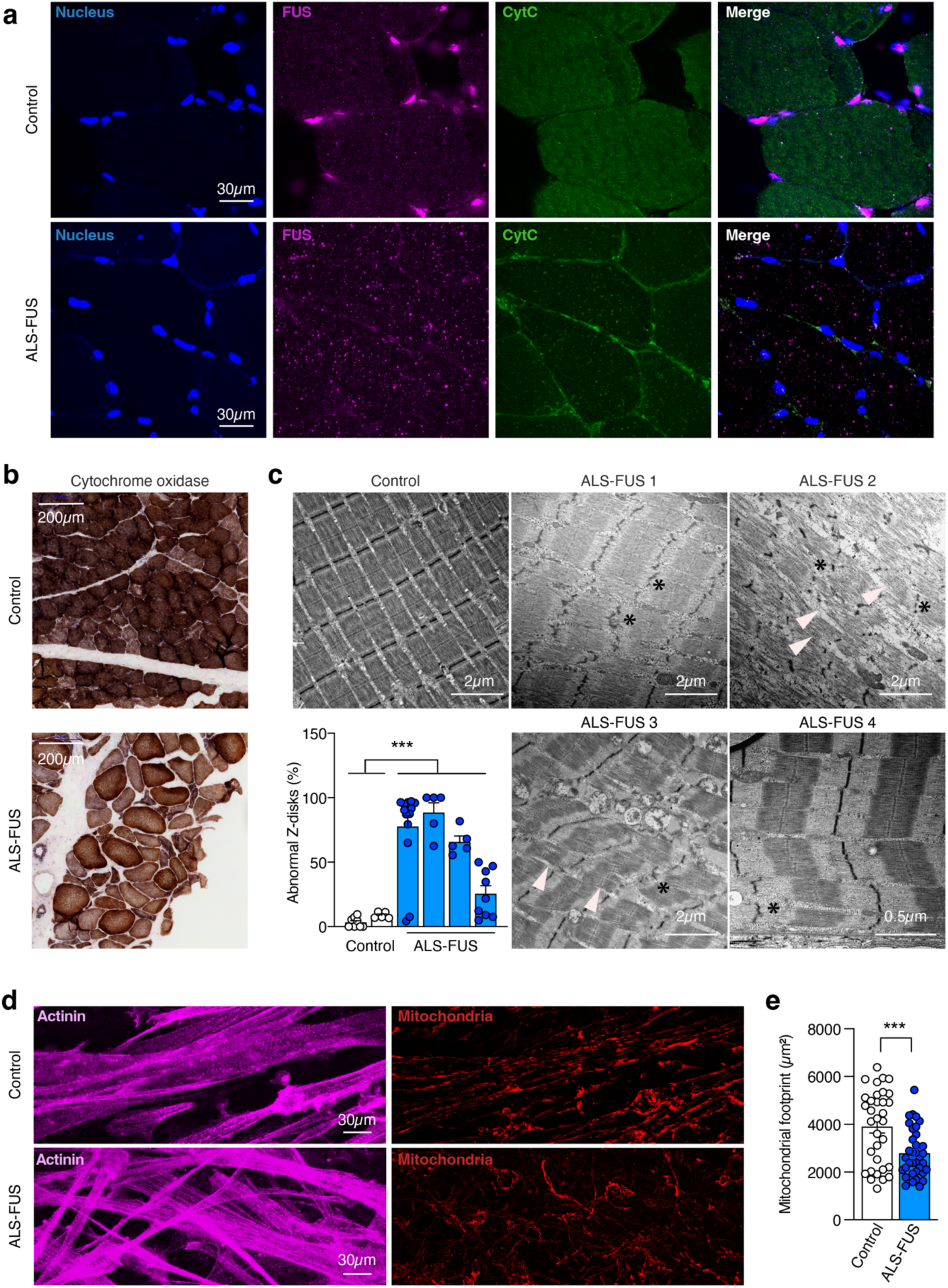
FUS mislocalization, abnormal sarcomeric ultrastructure, and mitochondrial defects in *FUS-*ALS muscles. a: Immunostaining for nucleus (DAPI, blue), FUS (magenta), and cytochrome c (CytC, green) on sections of muscle biopsies from a control individual (upper row) and a *FUS-*ALS patient (lower row). FUS protein was mainly localized to the nucleus in the control muscle, while cytoplasmic mislocalization was evident in *FUS-*ALS muscle. Cytochrome c staining was higher in control muscle compared to *FUS-*ALS muscle where it accumulated at the plasma membrane. Scale bar: 30 µm. b: Cytochrome oxidase histochemistry in control and *FUS-*ALS muscles. Scale bar: 200µm. c: Electron micrographs of muscles in healthy control showing normal ultrastructure with aligned Z-lines, and *FUS-*ALS patients showing extensive myofibrillar disorganization (white arrowheads), including disrupted Z-lines (stars). Lower left panel shows the percentage of fragmented, curvy and/or zigzag Z-lines in controls and *FUS-*ALS patients. Dots indicate the analysis of different images in the corresponding individuals. Nested t-test; ****p<0.0001. d-e: Representative mitochondrial morphology in spheres-derived myotubes from iPSCs of a *FUS-*ALS patient or its isogenic control. D: Cells were stained with Actinin (magenta) and Mitotracker (red). Scale bars: 30 µm. E: Quantification of mitochondrial footprint. Dots represent data from individual images obtained from 3 independent biological replicates. ***p<0.001, two-tailed unpaired t-test.

To exclude the possibility that these defects were secondary to muscle denervation during disease progression in the *FUS-*ALS patients, we studied the mitochondrial network in myotubes differentiated from a *FUS-*ALS patient iPSC line and its isogenic CRISPR-edited line where the *FUS* mutation was corrected ^33^. Control and *FUS-*ALS myotubes were stained with Mitotracker followed by immunofluorescence staining for Actinin. Disrupted mitochondrial networks were observed in *FUS-*ALS myotubes as compared to control myotubes (**Figure 1d**). Analysis of mitochondrial footprint (the total area covered by the mitochondrial network) indicated that *FUS* mutation leads to a significant decrease in mitochondrial network area in the *FUS-*ALS myotubes relative to control (**Figure 1e**). Mitochondrial ultrastructure, assessed using TEM, was not altered in *FUS-*ALS myotubes (**Supplementary** Figure 1b) but Z-lines were less dense in patient-derived myotubes compared to the isogenic control (**Supplementary** Figure 1b).

### Expression of mutant FUS in myocytes is required for muscle defects in mice

To investigate further the consequences of *Fus* mutation on muscle structure, we exploited two mouse models either knocked-out for the *Fus* gene (*Fus^-/-^*) or carrying a truncating mutation of FUS deleting the nuclear localization signal (NLS, *Fus^ΔNLS/ΔNLS^*). For both *Fus* alleles, homozygous mice die at birth of respiratory insufficiency ^25^. Therefore, we performed TEM on muscles at E18.5. Consistent with results in patient- or iPSC-derived samples, we observed profound abnormalities in sarcomere ultrastructure, including faint Z-lines and disorganized myofibrils in both *Fus^-/-^* and *Fus^ΔNLS/ΔNLS^* embryos (Figure 2a-b). These myofibrillar defects were accompanied by swollen mitochondria with loss of cristae (arrowheads in Figure 2a-b). Heterozygous *Fus^ΔNLS/+^* mice carrying one mutant *Fus* allele, similar to the genetic situation in *FUS*-ALS patients, reach adulthood and show adult-onset muscle weakness and progressive motor neuron degeneration ^24^.

**Figure 2:**
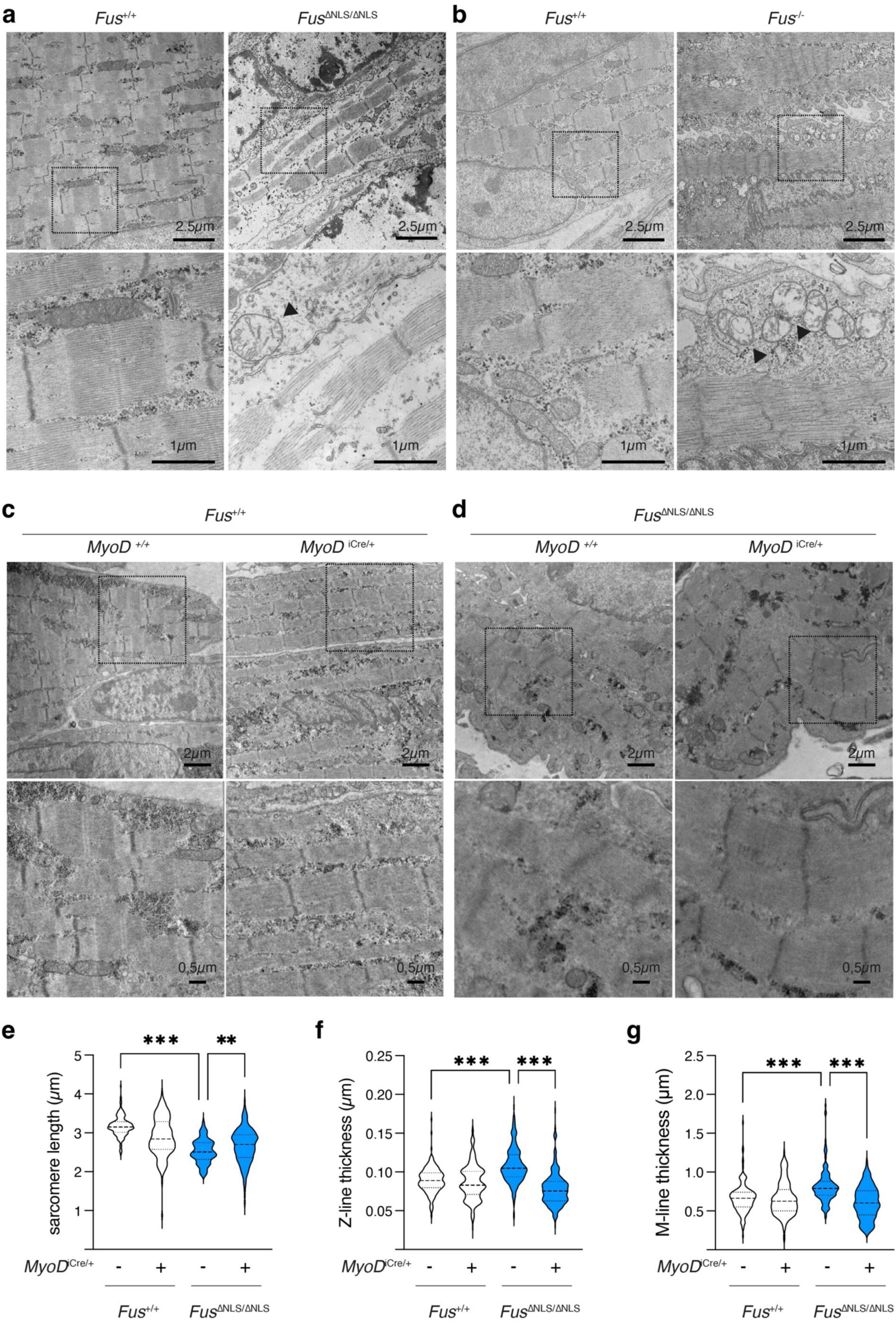
Homozygous *Fus* mutation or deletion in mice lead to muscle-autonomous ultrastructural defects a-b: Representative TEM images of gastrocnemius muscles of E18.5 *Fus^ΔNLS/ΔNLS^* (a), *Fus^-/-^* (b), and their respective control littermates. A higher magnification of the indicated regions (dashed boxes) is shown in the lower panels. Scale bars: 2.5 μm (upper panels) and 1 μm (lower panels). Arrowheads in lower panels indicate swollen mitochondria. c-d: Representative TEM images in gastrocnemius muscles of E18.5 *Fus^+/+^* (c) or *Fus^ΔNLS/ΔNLS^* (d), with or without MyoD^iCre/+^. A higher magnification of the indicated regions (dashed boxes) is shown in the lower panels. e-g: Quantitative analyses of ultrastructural parameters in sarcomeres of E18.5 *Fus^+/+^* or *Fus^ΔNLS/ΔNLS^*, with or without MyoD^iCre/+^. *Fus^+/+^*: 9 animals, 697 sarcomeres analyzed; *Fus^ΔNLS/ΔNLS^*: 10 animals, 755 sarcomeres analyzed; *Fus^+/+^*/MyoD^iCre/+^: 4 animals, 358 sarcomeres analyzed; *Fus^ΔNLS/ΔNLS^* / MyoD^iCre/+^: 3 animals, 276 sarcomeres analyzed. **P<0.01, ***P<0.001 by Kruskal Wallis test.

Notably, 22-month-old *Fus^ΔNLS/+^* mice also showed muscle ultrastructural defects, with swollen mitochondria and sarcomeric defects, in particular in M-lines and Z-lines, similar to homozygous E18.5 animals (**Supplementary** Figure 2a-e). Thus, ALS-related mutations in *FUS* lead to muscle structural defects, including disruption of sarcomeres and mitochondria, in both human and mouse.

The ΔNLS allele carries LoxP sites allowing CRE-dependent reversal of the mutation to wild-type *Fus* ^25^. To determine whether the observed muscle defects were attributable to intrinsic toxicity of mutant FUS in myocytes, we crossed *Fus^ΔNLS^* mice with *MyoD^iCre^* mice ^34,35^, allowing targeting of myocytes as early as E10.5, as previously described and validated ^30^. TEM analysis of muscles of *MyoD^iCre/+^; Fus^ΔNLS/ΔNLS^* showed a significant rescue of decreased sarcomere length and increased Z-line and M-line thickness in E18.5 muscle as compared to littermate *Fus^ΔNLS/ΔNLS^* mice (Figure 2c-g), demonstrating that these defects originated from expression of the mutant gene in myocytes. Thus, expression of mutant FUS in mouse skeletal muscle is required to trigger the ultrastructural defects.

### *Drosophila caz* function in myocytes is required for maintenance of adult muscle integrity

To evaluate whether the muscle cell-autonomous function of FUS is evolutionary conserved, we next turned to *Drosophila*. *Cabeza* (*caz*) is the single *Drosophila* orthologue of *FUS* and the two other FET family proteins EWSR1 and TAF15. We previously showed that *caz* is required for proper muscle development by promoting founder myoblast selection ^36^. Here, we evaluated a possible role for Caz in muscle maintenance by selectively inactivating *caz* in differentiating myocytes, after founder myoblast selection and myoblast fusion occurred. We crossed flies that selectively express FLP recombinase in myocytes (*MHC-GAL4>UAS-FLP*) to conditional *caz* knock-out animals in which the *caz* gene is flanked by FRT sites *(caz^FRT^)* ^37^. Because both full-body *caz* knock-out and myoblast-selective *caz* inactivation resulted in a reduced number and misorientation of adult abdominal muscles in pharate adult flies ^36^, we evaluated dorsal abdominal muscle morphology in flies in which *caz* was selectively inactivated in myocytes. Myocyte-selective *caz* inactivation resulted in a significantly reduced number of dorsal abdominal muscles, as well as occasional misorientation of these muscles in the abdominal body segments A3 and A4 (Figure 3a-d). Consistent with defective dorsal abdominal muscle morphology and the fact that abdominal muscle function is required for adult eclosion from the pupal case, myocyte-selective *caz* inactivation resulted in a lower-than-expected number of adult offspring, indicative of an eclosion defect (Figure 3e). To further evaluate the effect of myocyte-selective *caz* inactivation on adult muscle function, we determined the adult motor performance of ‘adult escaper’ flies using a negative geotaxis climbing assay ^38^ and identified a significant motor deficit (Figure 3f). In addition, the lifespan of myocyte-selective *caz* knock-out flies was significantly reduced (Figure 3g). Together, these results indicate that *Drosophila caz* function in myocytes – after myoblast fusion occurred – is required for structural and functional maintenance of adult muscles.

**Figure 3:**
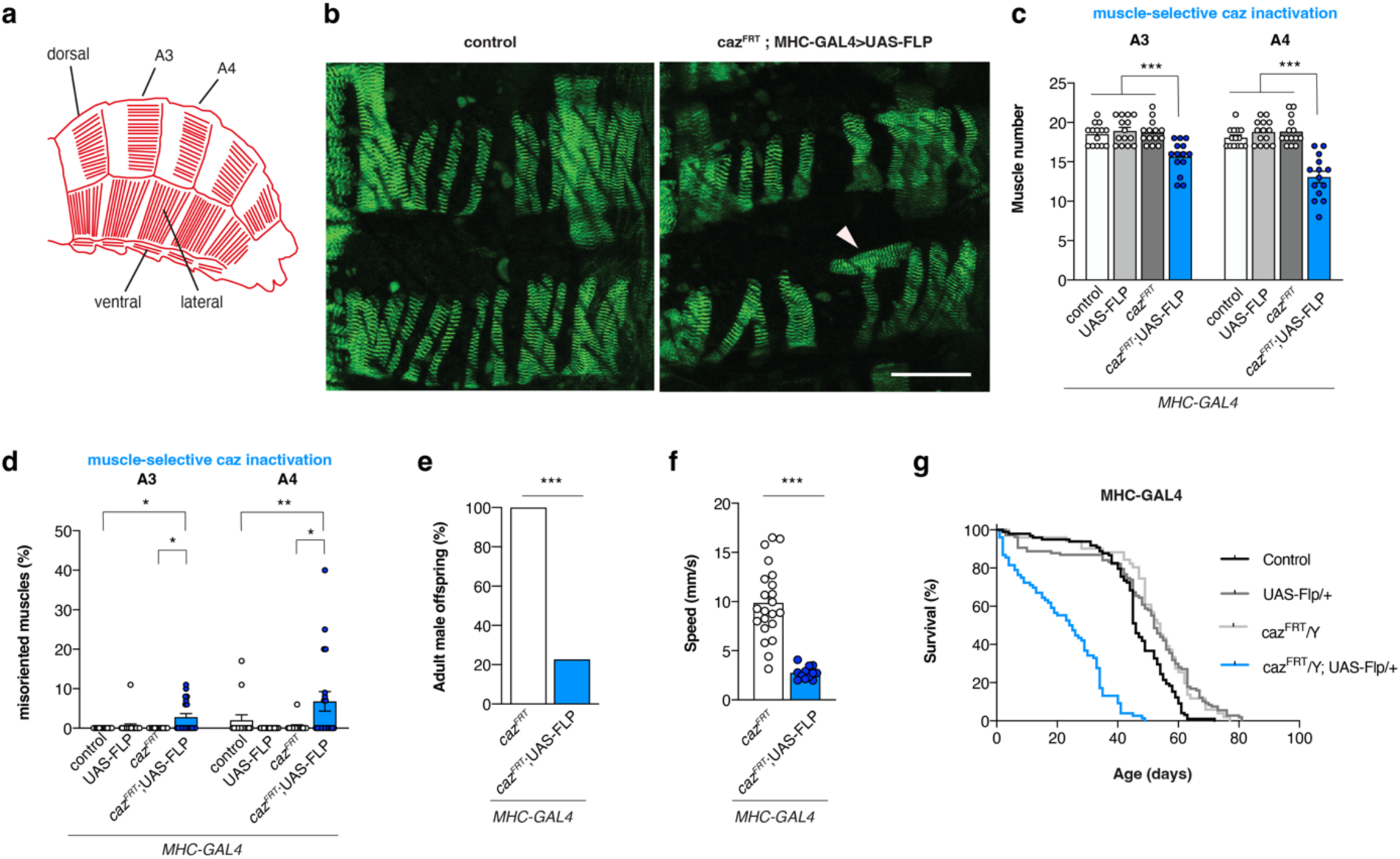
Selective *caz* inactivation in myocytes, starting after myoblast fusion, induces muscle defects, motor deficits and reduced life span. a: Schematic of adult abdominal muscle architecture in *Drosophila.* Dorsal, lateral and ventral abdominal muscles are indicated. b: Representative images of dorsal abdominal muscle defects in pharate adult flies (96h APF: 96 hours after pupa formation) in which *caz* was selectively inactivated in myocytes (*caz^FRT^*; *MHC-GAL4>UAS-FLP*) as compared to control animals. Arrowhead indicates a misoriented muscle. Scale bar: 100µm c-d: Quantification of the number (c) and misorientation (d, shown as % per animal) of dorsal abdominal muscles in abdominal segments A3 and A4 of 96h APF male pupae in which *caz* was selectively inactivated in myocytes (*caz^FRT^*; *MHC-GAL4*>UAS-FLP) as compared to control genotypes. (c) Two-way ANOVA with Tukey’s multiple comparisons test; ***p<0.0005; n=14 per genotype. Average ± SEM. (d) One sample Wilcoxon signed rank test to compare all genotypes to control, Mann-Whitney test to compare *caz^FRT^* to *caz^FRT^*; UAS-FLP; *p<0.05, **p<0.01; n=14 control, 20 UAS-FLP, 14 *caz^FRT^*, 20 *caz^FRT^*; UAS-FLP. e: Adult offspring frequency of male flies in which *caz* is selectively inactivated in muscle (*caz^FRT^*; *MHC-GAL4*>UAS-FLP) relative to *caz^FRT^*; *MHC-GAL4*/+ control flies (100%). Chi-square test; ***p<0.0001; n ≥ 214 F1 male flies per cross. f: Climbing speed (in mm/s) in an automated negative geotaxis assay of 7-day-old *caz^FRT^*; *MHC-GAL4*>UAS-FLP males as compared to *caz^FRT^*; *MHC-GAL4*/+ control males. Mann-Whitney test; ***p<0.0001; n=22 groups of 10 *caz^FRT^* flies versus 12 groups of 10 *caz^FRT^*; UAS-FLP flies. Average ± SEM. g: Life span of *caz^FRT^*; *MHC-GAL4*>UAS-FLP males as compared to the relevant control genotypes. Log-rank test; n=98 control, 107 UAS-FLP, 51 *caz^FRT^*, 76 *caz^FRT^*; UAS-FLP.

### FUS modulates the MEF2-dependent transcriptome in skeletal muscle

We then sought to delineate underlying molecular mechanisms of FUS-mediated muscle defects, and performed RNAseq on gastrocnemius muscles of E18.5 *Fus^-/-^* and *Fus^ΔNLS/ΔNLS^* mice and their respective littermate controls. We also included RNAseq analysis in C2C12 cells with or without si-RNA mediated knock-down of *Fus* (**Supplementary** Figure 3a). In all three models, we observed extensive transcriptional alterations (Figure 4a-c), with more than 3000 genes differentially expressed in each of the three models. Gene ontology (GO) analyses showed that downregulated genes in E18.5 *Fus^-/-^* and *Fus^ΔNLS/ΔNLS^* muscles and in C2C12 cells with FUS knock-down were related to muscle ultrastructure, including sarcolemma, sarcomere and sarcoplasmic reticulum, and mitochondrial function (Figure 4d-e), while there was no specific GO term enrichment in the upregulated genes. Illustrating this, we observed downregulation of multiple sarcomeric protein-encoding transcripts including actin (*Acta1*) and proteins of the Z-line (e.g. *Myoz1, Myoz2, Pdlim3* or *Actn3*) in all three models (Figure 4f).

**Figure 4:**
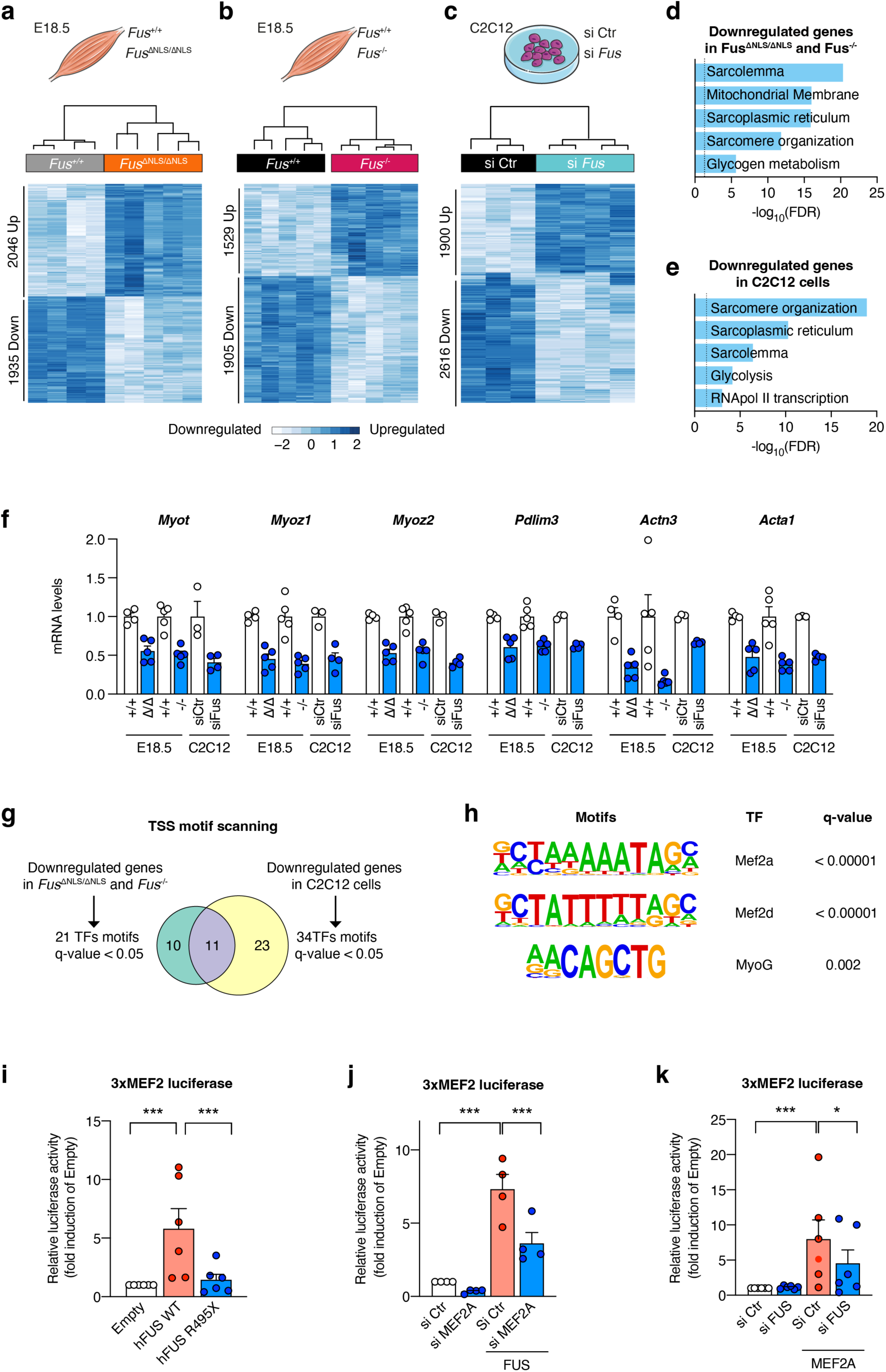
FUS modulates the MEF2-dependent transcriptome in muscle cells a-c: Heatmaps of differentially expressed genes in muscles from E18.5 *Fus^ΔNLS/ΔNLS^*mice (a), *Fus*^-/-^ mice (b) and their respective control littermates *Fus*^+/+^, or in mouse C2C12 myoblast cells (c) treated with siRNA against Fus (si Fus) or control (si Ctr). d-e: GO terms enriched in downregulated genes in *Fus*^-/-^ and *Fus^ΔNLS/ΔNLS^* muscles (d) or C2C12 cells (e). f: Normalized expression (based on FPKM from RNA-seq) of genes identified by RNA-seq to be significantly downregulated in *Fus*^-/-^ and *Fus^ΔNLS/ΔNLS^* muscles compared to their respective wild type controls or in C2C12 cells. Error bars represent SEM in 4-5 biological replicates. g: Venn diagram showing the overlap of transcription factor (TF) motifs enriched in the promoter region of downregulated genes in *Fus*^-/-^ and *Fus^ΔNLS/ΔNLS^* muscle and C2C12 cells with *Fus* knockdown. h: Enriched motifs in transcription start sites of downregulated genes predicted to be recognized by Mef2a, Mef2d and MyoG. i: Light units (RLU) relative to empty vector in C2C12 cells 24 h after transfection with plasmids expressing hFUS WT, hFUS R495X or an empty control plasmid, and a luciferase reporter plasmid carrying three MEF2 response elements controlling luciferase expression. Nested one-way ANOVA with Tukey’s multiple comparisons test; ***P < 0.0005, *P<0.05; N = 6 independent transfections. Each dot represents the mean of an individual experiment, each consisting of 4 technical replicates. j-k: Light units relative to siCtr in C2C12 cells 24 h after transfection of 3xMEF2 luciferase plasmid and siCtr (25 nM) or siMEF2A (25 nM, J) or siFus (25 nM, K), along with an expression plasmid for either hFUS_WT (J) or MEF2A (K) or empty control. Nested one-way ANOVA with Tukey’s multiple comparisons test ***P < 0.001, *P < 0.05. Each dot represents the mean of an individual experiment (n=4 for panel J; n=6 for panel K), each consisting of 8 (J) or 6 (K) technical replicates. Data represent the average ± SEM.

To determine whether alterations in a similar set of transcription factors could underlie the transcriptional changes in all three models, we performed motif scanning of the transcription start site (TSS) of downregulated or upregulated genes. While there was little enrichment in specific transcription factor motifs in upregulated genes, we identified 21 motifs bound by transcription factor enriched in downregulated genes in *Fus*^-/-^ and *Fus^ΔNLS/ΔNLS^* mice (q<0.05), and 34 in C2C12 cells with *Fus* knockdown. Of these transcription factor motifs, 11 were common to all three models (Figure 4g), with a striking enrichment in motifs recognized by MEF2 transcription factor family, in particular MEF2A and MEF2D (Figure 4h).

To more directly determine FUS may regulate MEF2-driven transcription in muscle cells, we transfected C2C12 cells with a reporter plasmid carrying luciferase under the control of three MEF2 response elements (3xMEF2-Luc). Co-transfection of a vector expressing wild-type FUS, but not a vector expressing ALS-linked R495X mutant FUS, was able to increase the activity of the 3xMEF2-luc reporter by 2- to 10-fold (Figure 4i). This was, at least in part, dependent upon endogenous MEF2A expression as FUS-driven 3xMEF2-luc reporter activity was blunted by simultaneous knockdown of MEF2A using siRNA (Figure 4j**, Supplementary** Figure 3b). Consistently, MEF2A overexpression increased 3xMEF2-luc activity, and this increase was reduced by knockdown of endogenous FUS (Figure 4k**, Supplementary** Figure 3a). Thus, FUS modulates MEF2-dependent transcription, which could underlie the observed muscle defects.

### FUS binds to and transactivates genes regulated by the ETS transcription factors of the PEA3 family

To determine whether the observed transcriptional alterations could be directly driven by FUS itself, we performed FUS ChIP-seq in C2C12 cells. We observed that FUS binds to relatively few genomic binding sites, with 294 significant peaks identified in C2C12 cells (Figure 5a). Peaks identified in FUS ChIP-seq but not in control IgG ChIP-seq were mostly located around transcription start sites of genes (Figure 5b-c). A typical FUS binding site is shown in Figure 5d close to the transcription start sites of the *Mrps18c* and *Helq* genes. FUS binding sites were enriched in genes involved in mitochondrial function, in particular mitochondrial translation (Figure 5e). Motif analysis of FUS binding peaks (Figure 5f) revealed a striking enrichment in a motif that is the predicted binding site of ETS transcription factors of the PEA3 family (ETV1, 4 and 5) (Figure 5g), as illustrated by the presence of 3 such PEA3-binding motifs in the illustrated *Mrps18c* promoter (Figure 5d).

**Figure 5:**
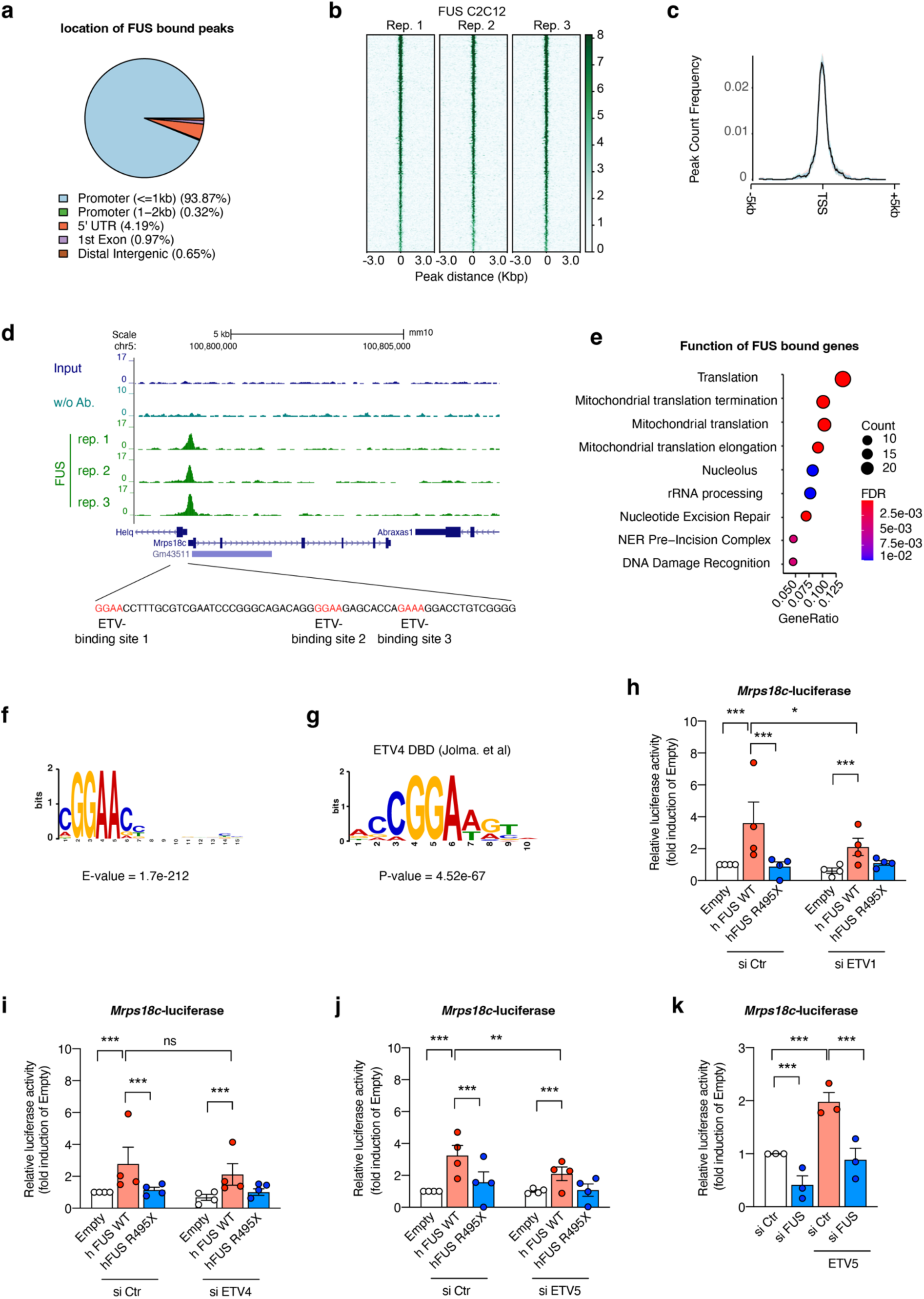
FUS is a transcriptional co-activator of PEA3 transcription factors a: Pie chart showing the distribution of FUS binding sites identified by ChIP-seq analysis in C2C12 cells. FUS binds mostly to promoters, close to the transcription start sites (TSS). b-c: Mean profiles established for FUS binding at TSS using SeqMiner in 3 independent replicates. d: Genome browser visualization of FUS binding on the promoter region of the *Mrps18c* and *Helq* genes. e: Gene ontology analysis of genes with FUS binding sites identified by ChIP-seq in C2C12 cells. f-g: MEME motif enriched in ChIP-seq peaks bound by FUS (F) and the predicted motif bound by PEA3 transcription factor ETV4 (G). h-j: Light units relative to empty control plasmid in C2C12 cells 24 h after transfection of an *Mrps18c-*luciferase plasmid and an expression plasmid for either hFUS WT, hFUS R495X or empty control. Cells were simultaneously transfected with si Ctr (25 nM) or si ETV1 (25 nM, H), si ETV4 (25 nM, I) and si ETV5 (25nM, J). Nested one-way ANOVA with Tukey’s multiple comparisons test; ***P < 0.001, **P<0.01, *P < 0.05. Each dot represents the mean of an individual experiment (n=3), each consisting of 4 technical replicates. Data represent the mean ± SEM. k: Light units relative to si Ctr in C2C12 cells 24 h after transfection of *Mrps18c-*luciferase plasmid and si Ctr (25 nM) or si FUS and an expression plasmid for either ETV5 or empty control. Nested one-way ANOVA with Tukey’s multiple comparisons test; ***P < 0.001. Each dot represents the mean of an individual experiment (n=3), each consisting of 4 technical replicates. Data represent the mean ± SEM.

To determine whether FUS could co-activate genes with ETS transcription factors, we generated a luciferase reporter under the control of the *Mrps18c* promoter that includes three ETS transcription factor motifs at the FUS binding site (Figure 5d). Wild type, but not R495X, FUS overexpression induced *Mrps18c* promoter activity (Figure 5h-j). Notably, all three PEA3 ETS transcription factors were expressed at significant levels in embryonic muscle and C2C12 cells (**Supplementary** Figure 4), suggesting that FUS action on its identified genomic binding sites could be dependent on either of the three PEA3 ETS transcription factors. Consistent with this notion, FUS mediated activation of *Mrps18c* promoter was reduced by simultaneous knock-down of *Etv1* or *Etv5,* but not *Etv4* (Figure 5h-j). Conversely, ETV5 overexpression increased *Mrps18c* promoter activity, and this was prevented by simultaneous FUS knock-down (Figure 5k). Together, these data suggest that FUS and PEA3 transcription factors cooperate to stimulate transcription of target genes.

### ETS transcription factors of the PEA3 family are required for muscle maintenance and reduced *Etv5* levels aggravate *FUS-*mediated muscle weakness in mice

Since FUS coactivates transcriptional activity of the PEA3 family transcription factors, we reasoned that loss of PEA3 family genes may phenocopy loss of FUS function. In *Drosophila*, *Ets96B* is the single ortholog of the mammalian *ETV1*, *ETV4*, and *ETV5* genes. Consistent with our hypothesis, myocyte-selective knock-down of *Ets96B* resulted in a reduced number of dorsal abdominal muscles for two independent transgenic RNAi lines (Figure 6a-b), a very similar phenotype as that induced by myocyte-selective *caz* inactivation (Figure 3). To extend this analogy in a mammalian system, we crossed *Etv5* knock-out mice ^39^ to *Fus*^ΔNLS/+^ mice. *Etv5^+/-^* mice did not show muscle weakness as evaluated using inverted grid test ^24^ (Figure 6c). However, heterozygosity of *Etv5* deletion triggered earlier onset and more severe muscle weakness in *Fus*^ΔNLS/+^ mice (Figure 6c). In summary, FUS and ETV5 are synergistically required to maintain muscle function through a mechanism conserved across species.

**Figure 6:**
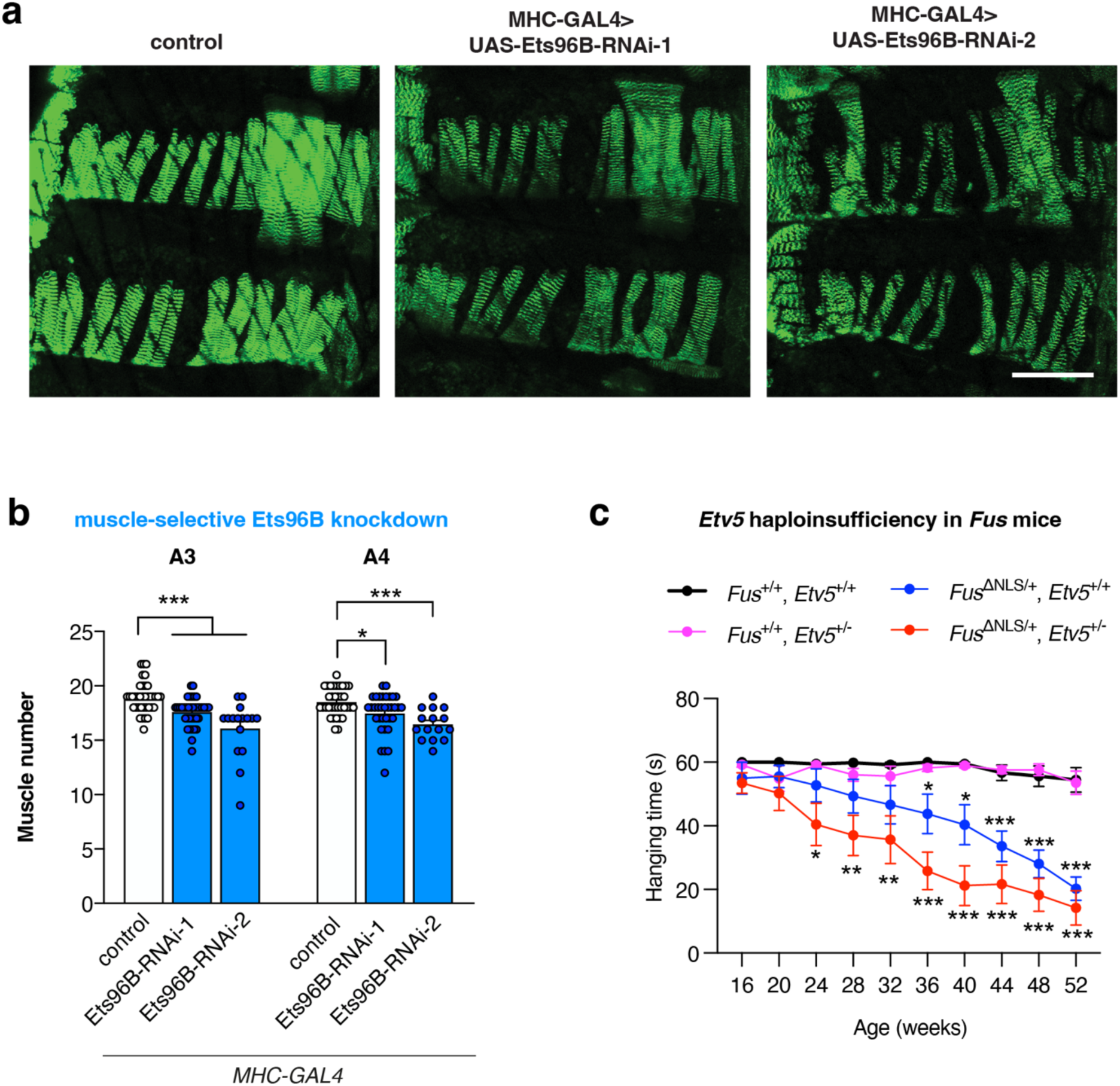
Reduced expression of *ETV5* orthologues induces muscle defects in *Drosophila* and enhances motor deficits of *Fus*^ΔNLS/+^ mice. a: Representative images of dorsal abdominal muscles in pharate adults (96h APF) in which Ets96B was selectively knocked-down in myocytes (MHC-GAL4>UAS-Ets96B-RNAi) using two independent RNAi lines as compared to control. Scale bar: 100µm. b: Quantification of the number of dorsal abdominal muscles in abdominal segments A3 and A4 of 96h APF male pupae in which Ets96B was selectively knocked-down in myocytes (as compared to control). Kruskal-Wallis test; *p<0.05, ***p<0.001; n=37 control, 37 Ets96B-RNAi-1, and 16 Ets96B-RNAi-2. c: Hanging time in the inverted grid test of male *Fus*^ΔNLS/+^ x *Etv5*^+/-^ mice from 16 to 52 weeks of age. n = 9 or 10 mice per genotype. *p<0.05, **p<0.01, ***p<0.005 by one-sample t-test with theoretical mean 60 s.

### FUS co-activates MEF2 and ETV5 dependent transcription

Our previous results showed a direct functional connection between FUS and PEA3 ETS transcription factors in muscle, and that *Fus* mutation hampers MEF2-dependent transcription. Notably, previous studies had suggested functional relationships between MEF2 and ETS transcription factors. Indeed, MEF2 co-activates the activity of ETV transcription factors during myogenic differentiation ^40^, and MEF2 and ETS transcription factors share genomic binding sites ^41^. Indeed, analysis of ChIP-seq datasets demonstrated that MEF2A and ETV5 binding sites were present at locations of FUS genomic binding (Figure 7a) with a significant overlap between MEF2A, ETV5 and FUS binding sites (Figure 7b). While only a small proportion of Etv5 and Mef2a peaks were also bound by FUS, almost all FUS binding sites were also bound by ETV5 and/or MEF2a (Figure 7b and **Supplementary** Figure 5). Consistently, gene set enrichment analysis (GSEA) showed significant enrichment between FUS peaks and genes downregulated following siRNA mediated knockdown of *Fus* in C2C12 cells and E18.5 Fus^ΔNLS^ muscles, and between MEF2A binding and genes downregulated in all three datasets (**Supplementary** Figure 6), suggesting a direct functional co-activation of MEF2A target genes by FUS and ETV5. Indeed, knockdown of ETV5, but not of ETV1 or ETV4, blunted activation of MEF2 transcriptional activity by FUS 24h after transfection of a MEF2 dependent luciferase reporter (Figure 7c-e). These data indicate that ETV5 is required for FUS-mediated co-activation of MEF2A target genes.

**Figure 7:**
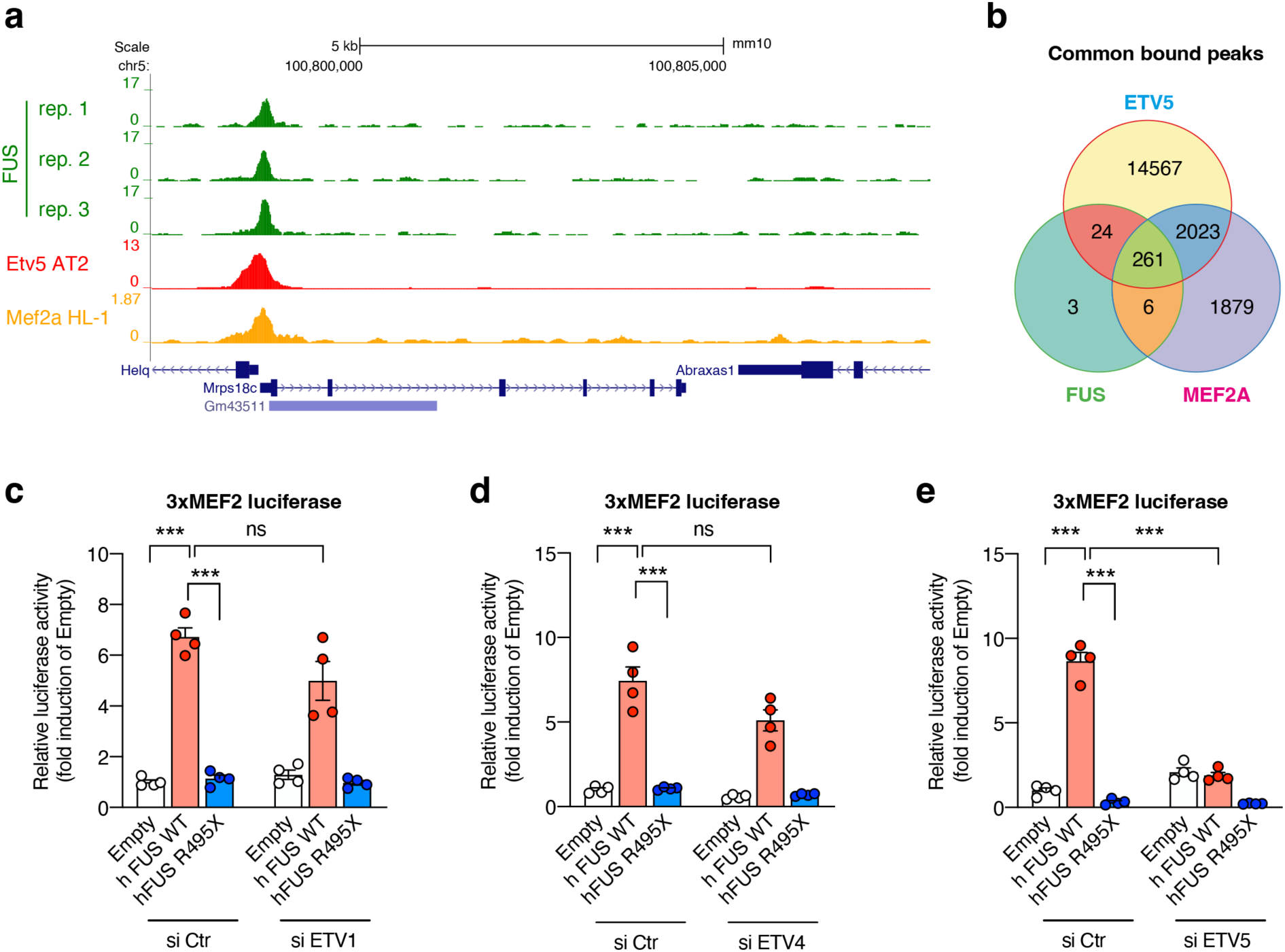
ETV5 is required for FUS-mediated co-activation of MEF2A target genes. **a:** Genome browser snapshot of MEF2A, ETV5 and FUS ChIP-seq binding sites at the *MRPS18C/HELQ* locus. b: Venn diagram showing the overlap between FUS, MEF2A and ETV5 binding sites. Hypergeometric p-values: Fus vs Etv5: 3.66×10^-248^; Fus vs Mef2a: <5×10^-324^; Etv5 vs Mef2a: <5×10^-324^. c-e: Light units relative to Empty vector + si Ctr in C2C12 cells 24 h after transfection of a *3xMEF2* luciferase plasmid and an expression plasmid for either hFUS WT, hFUS R495X or empty control. Cells were simultaneously transfected with si Ctr (25 nM) or si ETV1 (25 nM, C), si ETV4 (25 nM, D), and si ETV5 (25nM, E). ETV5, but not ETV1 or ETV4, is required for co-activation of the MEF2 reporter by FUS. Nested one-way ANOVA with Tukey’s multiple comparisons test; ***P < 0.001, **P<0.01, *P < 0.05. Each dot represents the mean of an individual experiment (n=4), each consisting of 4 technical replicates. Data represent the mean ± SEM.

### FUS phase separation promotes recruitment of MEF2A and ETV5 and is required for transactivation of target genes

To further examine whether this functional interaction could involve direct protein interaction, we produced fluorescent protein-tagged versions of MEF2A and ETV5 and performed co-immunoprecipitation of FUS using recombinant proteins. Both MEF2A and ETV5 proteins interacted with FUS alone, and MEF2A/FUS interaction was increased by addition of ETV5 (Figure 8a-b). Electrophoretic mobility shift assays (EMSA) using a *Mrpl30*-derived DNA probe encompassing the FUS binding motif at this gene showed that addition of FUS and MEF2A increased the binding affinity of ETV5 to its target probe (Figure 8c-d). Similar results were obtained for ETV1, but not ETV4 (**Supplementary** Figure 7a-b). Importantly, mutating the ETV5 binding motifs in the probe abolished binding (**Supplementary** Figure 7c-d).

**Figure 8:**
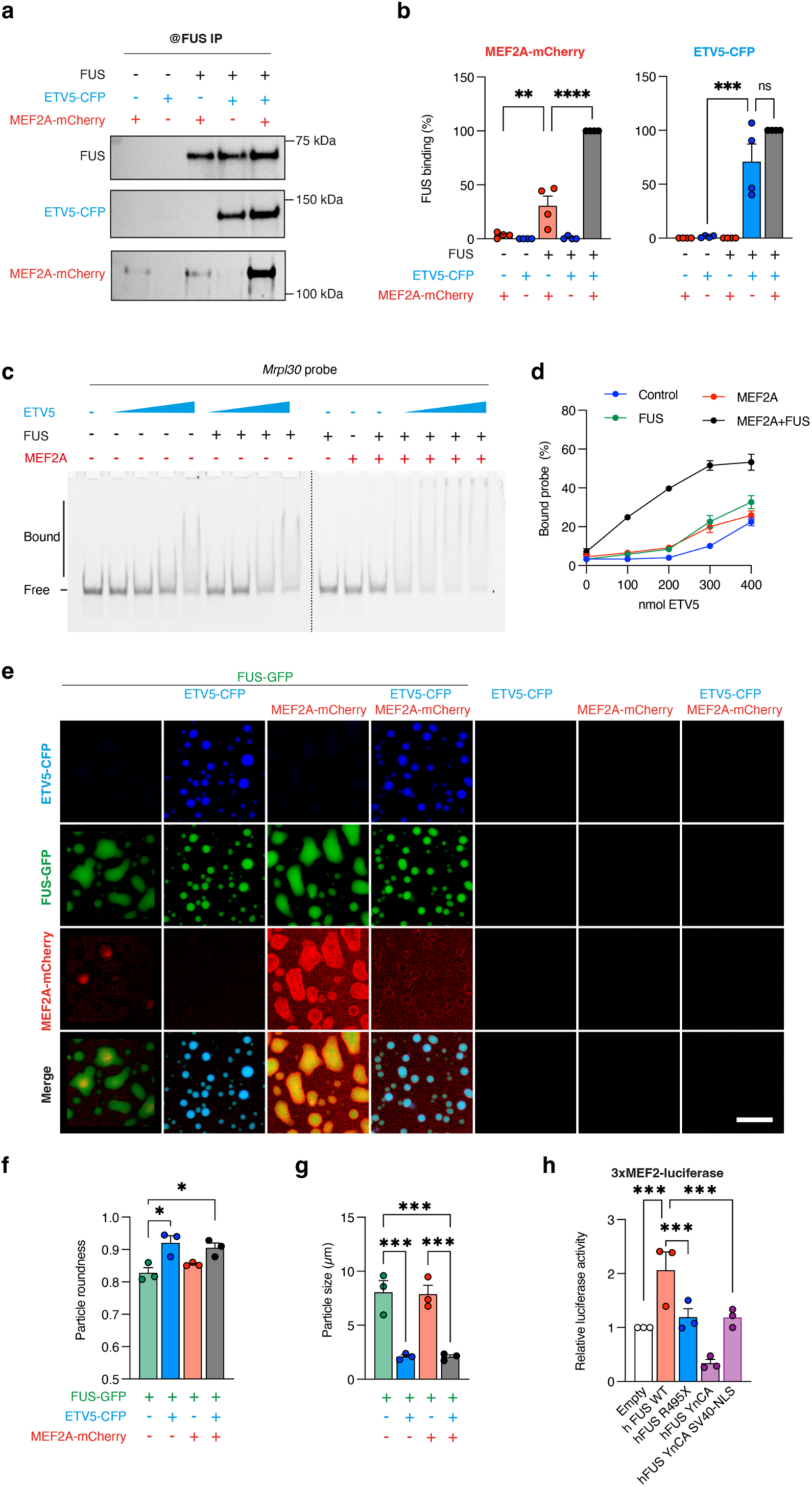
FUS binds to and phase separates MEF2A and ETV5 and allows cooperative binding on DNA target sites. **a-b:** Recombinant FUS protein was incubated in presence of ETV5-CFP or/and MEF2A-mCherry recombinant proteins. After immunoprecipitation using FUS antibody then pulled out with A/G-coupled magnetic beads, immunoprecipitated proteins were examined by western blot using FUS, ETV5 and MEF2A antibody. The band intensities of co-immunoprecipitated MEF2A-mCherry and ETV5-CFP were quantified using iBright Analysis Software (B). Error bars represent SEM of four independent experiments. c-d Cooperative binding assay of FUS, ETV5, MEF2A recombinant proteins to FAM labelled *Mrpl30* promoter-derived DNA probe. EMSA experiments performed by using increasing concentrations of ETV5 (60, 120, 180 and 240 nM) and constant concentration of FUS (300 nM) and/or MEF2A (50 nM). Panel D shows the percentage of *Mrpl30* probe bound across 3 independent experiments. The intensity of the signals of the protein-*Mrpl30* probe complex was quantified using the iBright Analysis Software. Error bars represent SEM of three independent experiments. e: Representative confocal images of FUS-EGFP condensates (5 µM) in absence or presence of ETV5-CFP (3 µM) and/or MEF2A-mCherry (0.85 µM) upon protease-mediated cleavage of the MBP-solubility tags. No condensates were observed for ETV5 or MEF2A alone or mixed together in absence of FUS condensates. Bar, 10 µm. f-g: Quantification of FUS-EGFP condensate roundness (F) and size (G). Mean ± SEM of three independent replicates with at least 179 condensates in total from ≥3 different fields of view analyzed. * p<0.05, *** p<0.005 by one-way ANOVA with Tukey’s multiple comparison test. h: Light units relative to Empty vector in C2C12 cells 24 h after transfection of *3xMEF2* luciferase plasmid and an expression plasmid for either hFUS WT, hFUS R495X, LLPS deficient hFUS (hFUS YnCA), LLPS-deficient hFUS with SV40 NLS or empty control plasmid. Nested one-way ANOVA with Tukey’s multiple comparisons test; ***P < 0.001, **P<0.01, *P < 0.05. Each dot represents the mean of an individual experiment (n=3), each consisting of 4 technical replicates. Data represent the mean ± SEM. Expression of human wild type FUS but neither R495X nor LLPS deficient FUS increases activity of the MEF2 reporter.

FUS is known to undergo liquid-liquid phase separation (LLPS) at physiological concentrations ^42^ and LLPS is critically involved in chromatin compartmentalization during transcription ^43^ and other gene regulatory processes ^44^. We thus hypothesized that FUS LLPS could be critical in mediating ETV5 and/or MEF2 transcriptional function. Interestingly, both ETV5 and MEF2A are enriched in intrinsically disordered regions, as predicted using PONDR ^45^ (**Supplementary** Figure 8a) and were predicted to undergo phase separation using PSP predictor ^46^ (**Supplementary** Figure 8a). Despite these predictions, neither ETV5-CFP nor MEF2A-mCherry or the combination of both proteins were able to form condensates on their own, at least at low µM concentrations after 60 minutes (Figure 8e). To assess whether presence of FUS might trigger LLPS of ETV5 and/or MEF2A, we introduced MBP-FUS-GFP ^42^, whose LLPS can be induced by Tev protease-mediated cleavage of the MBP solubility tag. When FUS LLPS was triggered in presence of ETV5-CFP or MEF2A-mCherry, either protein partitioned into FUS condensates, suggesting that ETV5 and MEF2A might get recruited and concentrated at promotors through FUS condensation. While MEF2A did not affect FUS droplet formation and morphology, presence of ETV5 led to smaller and rounder FUS droplets, indicative of less droplet fusion and reduced droplet dynamicity (Figure 8e-g). Abrogation of LLPS properties of FUS, through the mutation of 7 tyrosine residues to alanine ^47^ abrogated FUS-mediated activation of 3xMEF2-luc reporter activity, even after addition of a SV40 NLS to force entry of the LLPS deficient FUS in the nucleus (Figure 8h). Thus we propose that LLPS properties of FUS may help to recruit MEF2A and ETV5 and is required for FUS-mediated activation of these transcription factors.

## Discussion

Our study identifies a critical function of FUS in muscle development and maintenance through co-activation of key transcription factors involved in muscle differentiation. Importantly, we also provide evidence that FUS mutations in ALS patients lead to similar muscle defects. We further show that two of these pathways synergize through FUS-mediated phase separation of ETV5 and MEF2. These findings have major implications for our understanding of physiological functions of FUS, and to inform the design of novel therapeutic approaches for ALS-*FUS*.

Our results demonstrate that FUS is critical in muscle development and maintenance in multiple species. In mice, ablation of *Fus* or deletion of its nuclear localization sequence severely hampers muscle structure leading to severe defects in sarcomeric and mitochondrial ultrastructure. In Drosophila, deletion of the *Fus* ortholog leads to similar alterations in muscle development ^36^ and maintenance (this study). Furthermore, mutation of one allele in *FUS*, either in *FUS*-ALS patients or in *Fus^ΔNLS/+^* mice, significantly compromised adult muscle ultrastructure. Importantly, we provide several complementary lines of evidence in iPSC derived-models, mouse models and Drosophila models that the observed muscle phenotype is primarily caused by loss or altered expression of FUS in myocytes. This novel role for FUS in muscle development and maintenance adds on to the increasing evidence that mutations in ALS-associated genes may also lead to prominent muscle phenotypes. Indeed, mutations in *MATRN3* ^48^, *HNRNPA1* ^49^ or *CHCHD10* ^50^ lead to phenotypes ranging from pure myopathic syndromes to complex mitochondrial phenotypes and multi-system proteinopathy ^4,51^. Mutations in *TARDBP*, encoding TDP-43, have also been described in patients with myopathy ^52,53^ which may result from disruption of TDP-43’s function in RNA granules involved in muscle development and regeneration ^54^. To our knowledge, mutations in *FUS* in patients with a primary myopathic phenotype have not yet been identified, although reports exists of clinical myopathic features in FUS-ALS patients ^55^ and of altered neurophysiological or histological muscle properties in FUS-ALS patients, suggesting either initial or accompanying myopathy in these patients ^30,32,56^.

Our study identifies two different transcription factors interacting with FUS, which could contribute to the muscle phenotype. First, expression of genes controlled by MEF2 transcription factors, in particular MEF2A, is severely decreased in three models of FUS depletion or mutation. Interestingly, muscle defects observed upon *Fus* loss or mutation, are not recapitulated by single ablation of either *Mef2a* or *Mef2d* ^57^, yet *Mef2a* ablation leads to a similar developmental defect in cardiac muscle ^58^. It is possible that FUS is required for the transcriptional activity of both MEF2A and MEF2D, and that loss of FUS leads to functional loss of both MEF2 isoforms and a more drastic phenotype than loss of the single MEF2 isoforms. Alternatively, other transcription factors, such as *Mef2c* or *Myog* may also require FUS, leading to a more severe muscle phenotype in *Fus* mutant mice than in mice with a single mutation of any of these transcription factors.

Notably, we did not observe a significant enrichment of FUS on MEF2 genomic binding sites, while FUS binding was very specifically associated with *PEA3* transcription factor binding sites. A possible explanation is that FUS may only indirectly associate to its chromatin binding sites through binding of transcription factors. It is possible that the physical interaction between ETV5 and FUS is stronger than the looser FUS/MEF2 interactions, leading to better preservation through chromatin immunoprecipitation steps. Our observation of FUS binding to a subset of *PEA3* transcription factor binding sites in muscle cells is consistent with our recent results in hippocampus ^29^, and is also in line with our previous observation that FUS transactivates subsynaptic genes in an ERM/ETV5-dependent manner ^30^. Despite the binding of FUS to *PEA3* transcription factor target genes, the transcriptional outcome of FUS loss or mutation on these specific genes remains modest. We hypothesize that loss of FUS is compensated efficiently by other FET proteins, such as EWSR1 or TAF15. It is also possible that a strong transcriptional response induced by FUS requires MEF2 binding. Consistent with this hypothesis, downregulation of genes bound by FUS was much stronger when MEF2A was also binding to these genes, suggesting that MEF2A dependency was critical in driving the transcriptional defect upon loss of FUS. Importantly, our results show that MEF2 and *PEA3* transcription factor pathways intersect.

We found that FUS, MEF2A and ETV5 physically interact *in vitro*, and cooperatively bind to FUS DNA binding sites. Furthermore, both ETV5 and MEF2A partition into FUS condensates. While ETV5 potently modified FUS condensates morphology and appeared to make them smaller, possibly more rigid, MEF2A did not modify the condensate shape. As a result, the combination of FUS, ETV5 and MEF2A showed an accumulation of MEF2A on the FUS-ETV5 condensate surface. It seems possible that FUS and ETV5 thereby potentiate MEF2A transcriptional response through modification of condensate dynamics. Further research is needed to elucidate the relationships between these 3 proteins and their LLPS behavior in transcriptional regulation of muscle development. FUS, MEF2 and *PEA3* transcription factors are broadly expressed, and our findings could thus have consequences beyond muscle development.

In adult muscle, both *Fus* and *Mef2a* are expressed in myonuclei, with an enrichment in subsynaptic nuclei ^30,59,60^, while *Etv5* expression is restricted to subsynaptic nuclei ^61^. It is thus likely that the FUS/MEF2A/ETV5 pathway is more relevant for regulating gene expression in subsynaptic than extrasynaptic nuclei in adults, consistent with our previous study ^30^. Beyond skeletal muscle, both MEF2 and *PEA3* transcription factors are expressed in the central nervous system and have been involved in synaptic plasticity ^62^ or neuronal specification ^63^. It will be important to determine whether FUS also coactivates MEF2 and *PEA3* transcription factors in neurons.

Here we show that loss of FUS nuclear function triggers cell autonomous detrimental effects on muscles. This evidence has an immediate translational importance as current strategies to treat *FUS*-ALS are based on antisense oligonucleotides (ASOs) to reduce FUS expression ^64^. The rationale for lowering strategies in *FUS-*ALS is supported by several studies, including ours, showing that FUS mutation leads to dominant gain of function responsible for motor neuron loss ^24,25,27,28,64^. Intrathecal administration of non-allele specific ASOs targeting FUS is currently tested in *FUS-*ALS patients ^65^. Considering our observation that loss of FUS in skeletal muscle has deleterious consequences, it will be important to evaluate possible muscle damage as an adverse effect of chronic FUS downregulation.

In summary, we show here that FUS is critical for muscle development and maintenance, and provide evidence that this occurs, at least in part, through co-activation of two transcription factors, MEF2A and ETV5. The resulting muscle defects may also contribute to the FUS-ALS phenotype in patients.

## Methods

### Patient biopsies

#### Human skeletal muscle biopsies

For human skeletal biopsy samples, muscle specimens were extracted in an open biopsy of the triceps (FUS1), deltoid (FUS2), vastus lateralis (FUS3), and biceps (FUS4, CNTRLs 1-2) and prepared for cryosection or electron microscopy (**Supplementary Table 1**). Control subjects who presented at our center with chronic muscle pain, showed no myopathic changes in the biopsy results. Skeletal muscle was obtained for diagnostic reasons as a part of the regular clinical work-up, including electromyography and muscle biopsy. This study was conducted according to the Helsinki Declaration. All included patients gave informed written consent for scientific use of remaining biopsy material, which was approved by the local University of Ulm Ethics Committee of the University of Ulm (reference no. 20/10).

#### Immunohistochemistry and histology on patient muscle biopsies

Samples were frozen with chilled isopentane and stored at −80 °C. Serial cuts of 10 μm thick were made using a cryostat (Leica CM 3050S) and placed on Superfrost slides (VWR, 631-0448). Slides were dried for 20 minutes then fixed with 4% PFA in 0.1 mM sucrose for 15 minutes at room temperature. Sections were used for immunohistochemistry. After washing with PBS, sections were then permeabilized with 0.1% Triton X-100, and nonspecific binding sites were blocked with 10% goat serum and 5% FBS in PBS for 2 hours at room temperature. Rabbit anti-FUS (Sigma, HPA008784, 1:1,000) and mouse anti-Cytochrome c (BD Biosciences, 556432, 1:1000) were incubated overnight at 4 °C. After rinsing in PBS, Alexa Fluor 488 goat anti-rabbit (A-11034) and Alexa Fluor 647 goat anti-mouse (A-21235) (all 1:1000, Invitrogen) were incubated for 2 h at room temperature. Sections were then washed with PBS and water and mounted with ProLong Gold Antifade reagent with DAPI (Invitrogen, P36935).

#### Imaging of human tissues

All fluorescence images were acquired using a Leica TCS SPE II high-resolution spectral confocal microscope with a resolution of 1024 x 1024 pixels. The mean fluorescence intensity of cytochrome c was quantified in human muscle sections. Briefly, five random images were taken for each control and mutant section, followed by selecting regions of interest (ROIs) in each image. To select ROIs, transverse sections of the muscle were specifically chosen, and then ROIs were always placed inside the muscle fiber. Afterwards, mean fluorescence intensity for five ROIs was measured in each picture after applying the same settings to all samples using Image J Fiji software. Sections with Cytochrome c oxidase (COX) histochemical labeling were imaged using the MIRAX scan.

TEM images were acquired using a JEOL 1400 Transmission Electron Microscope (TEM), and the images were digitally recorded with a Veleta camera (Olympus). Quantification of TEM images was performed using Image J Fiji software. In human muscle sections, five images were randomly taken for longitudinal parts of the muscle with the same magnification for each control and patient sample. Aligned Z-disk were counted as normal Z-disks, while fragmented, zigzag, and curvy ones were counted as abnormal Z-disks. The percentage of abnormal Z-disks was determined for each image of the control and mutant samples. Regarding muscle mitochondria, the area of nearly 100 muscle mitochondria from random images of healthy control and FUS patient was measured using Image J.

### Human iPSCs culture and differentiation

#### Human iPSCs

The *FUS* and isogenic control hiPSC lines have been previously described and characterized^33^. Briefly, the *FUS* hiPSC line was derived from a 26 years old patient carrying a *de novo* frameshift mutation in *FUS (*R495QfsX527 (c.1483delC). The isogenic control cell line (Corrected^FUS^) was generated by inserting a cytosine in position 1483 and correcting the mutation using Clustered Regularly Interspaced Short Palindromic Repeats (CRISPR) technology. HiPSCs were cultured at 37 °C (5% CO2, 5% O2) on Matrigel®-coated (Corning, 354,277) 6-well plates using mTeSR1 medium (Stem Cell Technologies, 83,850). At 80% confluence, the colonies were detached using Dispase (Stem Cell Technologies, 07923) and passaged in a 1:3 or 1:6 split ratio.

#### Differentiation of myotubes

Differentiation of myogenic cells from human iPSCs was performed as previously published using a sphere-based culture method ^66^. Briefly, hiPSCs were cultured in Stemline medium (Sigma, SA3194) containing 100 ng/ml FGF-2, 100 ng/ml EGF, and 5 ng/ml heparan sulfate (Sigma, H7640-1mg) in suspension using ultra-low attachment flasks. Cells were split on a weekly basis using a 2 minutes accutase (Sigma, A6964) digestion, and media were exchanged every second day. After 6 weeks in culture, cells were dissociated using accutase digestion, and filtered through a 30 μm pre-separation filter (Miltenyi Biotec, 130-041-407). Then a total of 250,000 cells per well were seeded on PLO and laminin coated glass coverslips in 24-well plates. After 8 weeks of differentiation, mature myotubes were obtained and either fixed or lysed for analysis.

#### Mitochondrial network labeling and analysis

Mitochondrial labeling was carried out as previously described ^67^. In summary, derived myotube mitochondria were labeled with MitoTracker™ orange CMTMRos (50 nM final concentration, Thermofisher # M7510) for 40 min at 37 °C in muscle differentiation medium. Myotubes cultures were then fixed in 4% PFA for 15 minutes at room temperature (RT) and imaged. Mitochondrial networks were analyzed using the “MiNA plugin” in Image J which allows the semiautomated analysis of the mitochondrial network, providing a topological skeleton of mitochondria ^68,69^. Briefly, five images were acquired for the mitochondrial network of control and FUS-mutated-derived myotubes. Particle analysis took place after applying a threshold over the network, allowing measurement of the mitochondrial footprint. All experiments were performed in a minimum of N= 3 independent replicates (independent differentiations).

#### Immunocytochemistry in sphere-derived myotubes

Immunostaining was performed as previously described ^30^. Myotube cultures were fixed in 4% (Merck) and 10% sucrose (Roth, 4621.1) in PBS (DPBS, Gibco, H15-001) for 15 minutes at room temperature. After washing, samples were permeabilized with 0.2% Triton X-100 (Roche, 10789704001) for 10 minutes. After blocking with 5% FBS (Gibco, 10500) and 10% goat serum (Millipore, S26-100mL) in DPBS for 1 h, primary antibodies rabbit anti-FUS (Sigma, HPA008784, 1:1000) and mouse anti-actinin (Sigma, A7811, 1:500) were incubated for 48 h at 4 °C. After subsequent washing, the following secondary antibodies were incubated for 1 h at room temperature: Alexa Fluor 488 goat anti-rabbit (A-11034), Alexa Fluor 647 goat anti-mouse (A-21235) (all 1:1000, Invitrogen). Then samples were mounted using ProLong Gold Antifade reagent with DAPI (Invitrogen, P36935).

### Mouse husbandry and behavior

#### Mouse housing and breeding

Mice were housed in the Faculty of Medicine from Strasbourg University, the animal facility of the Max Planck Institute for Molecular Biomedicine, and the animal facility of Radboud University, with 12/12 hours of light/dark cycle and unrestricted access to food and water.

*Fus*^ΔNLS/+^ knock-in mice were generated in the Institut Clinique de la Souris (ICS, Illkirch, Strasbourg) and were maintained and genotyped as previously described ^24,25,30^. This mouse strain expresses a truncated FUS protein that lacks the PY-NLS, which is encoded by exon 15 of the *Fus* gene. This mutation can be reverted to re-express a wild type FUS protein upon CRE-mediated recombination ^24,25^. Mice heterozygous and homozygous for the targeted allele will hereafter be referred to as *Fus*^ΔNLS/+^ and *Fus*^ΔNLS/ΔNLS^, respectively.

*MyoD^iCre/+^* (FVB.Cg-Myod1tm2.1(icre)Glh/J; Strain #:014140^34,35^) and *Erm^-/+^* (Stock Etv5tm1Kmm/J, Strain #:022300) ^39^ mice were purchased from the Jackson Laboratory and maintained in the Animal Research Facility (Centraal Dierenlaboratorium) in Nijmegen. Animal experiments performed in Nijmegen were approved by the by the national Dutch ethics committee ‘Centrale Commissie Dierproeven’ (Centrale Commissie Dierproeven) under reference number AVD10300 2019 8544.

*Fus* knock-out (*Fus^-/-^)* mice were generated by the transgenesis facility of the Max Planck Institute for Molecular Biomedicine, using ES cells obtained from the European Conditional Mouse Mutagenesis Consortium (EUCOMM) and were previously described ^24,25^. The genetic background of mice used in this study is C57Bl/6J. *Fus*^ΔNLS/ΔNLS^*; MyoD^iCre/+^* mice have a C57Bl/6N background and *Fus*^ΔNLS/+^*; Erm^-/+^* have a mixed 50% C57Bl/6N, 50% 129S6/SvEvTac background.

#### Mouse motor behavior

The inverted grid test was performed every 4 weeks from 16 weeks of age until 52 weeks of age. Prior to testing, mice were habituated in the testing room for 30 min. Mice were allowed to hang on an inverted grid for up to 60 sec, and the hanging time was recorded and documented. The hanging time of each mouse was evaluated twice per session with 15 min breaks between tests.

### Drosophila experiments

Flies were kept in a temperature-controlled incubator with 12 hours on/off light cycle at 25°C. To induce myocyte-specific *caz* inactivation, *caz^FR^*^T^; MHC-GAL4,UAS-gma-GFP/Tm6b,Tb females were crossed to UAS-FLP (BDSC stock 4539) males and non-Tubby male offsprings were used for the experiments. For Ets96B knockdown experiments, UAS-Ets96B-RNAi-1 (BDSC stock 34783) and UAS-Ets96B-RNAi-2 (BDSC stock 31935) were crossed to MHC-Gal4, UAS-gma-GFP.

#### Imaging of dorsal abdominal muscles

To image dorsal abdominal muscles, male larvae were selected and placed in a new vial containing standard *Drosophila* medium supplemented with yeast paste. White prepupae were collected on wet Whatman filter paper in petri dishes and kept at 25°C for 96 h. After 96 h, pharate adults were dissected out from the pupal case and placed in an approximately 2 mm furrow of a plastic slide with a drop of 60% glycerol. Images were acquired using a Leica SP8 laser scanning confocal microscope. For quantifications, dorsal abdominal muscles (DAMs) were counted using Fiji (http://fiji.sc/Fiji) by going through Z-stacks.

#### Motor performance assay

To evaluate motor performance, male flies were collected within 48 h after eclosion and housed in groups of 10 individuals. Motor performance of 7-day-old flies was evaluated as described previously ^38^. Average climbing speed (mm/s) was determined and compared between genotypes.

#### Adult offspring frequency and life span analysis

To determine adult offspring frequencies, the number of adult flies per genotype, eclosing from relevant crosses were counted daily for 9 days. For life span analysis, male flies were collected on the day of eclosion and housed at a density of 10 flies per vial. The number of dead flies was counted every day and flies were transferred to fresh food vials every 2-3 days.

### C2C12 myoblast culture experiments

#### C2C12 myoblast culture

C2C12 myoblast cells were purchased from ATCC (ATCC® CRL-1772™). Cells were cultured in Dulbecco’s modified Eagle’s medium containing 10% Fetal Bovine Serum (Fisher scientific, 11531831) and 1% Penicilin-Streptomycin (Sigma, P4333) at 37 °C in an incubator with 5% CO_2_. Between 60 and 80% of confluency, myoblasts were differentiated into myotubes with differentiation medium (identical medium, with 0.1% Fetal Bovine Serum). Culture medium was changed every day and transfections were performed between 5-15 passages.

#### siRNA Transfection

C2C12 were cultured in 6-well plates until 60% of confluency. Cells were transfected with siRNA in differentiation medium using Lipofectamine RNA iMAX (Fischer scientific sas, 13778150) according to the manufacturer’s instructions. siRNA against *Fus*, *Mef2a*, *Etv1*, *Etv4*, *Etv5* and negative control siRNA were provided by Dharmacon. Cells were harvested 48h after transfection.

#### Plasmid Transfection and luciferase assay

C2C12 were cultured in 24 well-plates until 80% of confluency. Transfections were performed in differentiation medium with expression and reporter plasmids using TransIT-X2 (MIR6000, Myrus) according to the manufacturer’s instructions.

Expression vectors used for transfections were: pCMV empty plasmid, pCMV-Myc-FUS (expressing N-terminal myc-tagged human wild type FUS), pCMV-Myc-FUS-R495X, pCGN-MEF2A α2β ^70^. pCMV-MycFUS YnCA and pCMV-MycFUS YnCA-SV40 NLS were synthesized by Thermo scientific using sequences published by Qamar and collaborators ^47^. Reporter plasmids used are pGL3-MEF2-luc (termed 3xMEF2-luciferase, addgene, 32967) and *Mrps18C*-luciferase (Gene Copoeia). After 24h of transfection, proteins were extracted and luciferase activity was measured (Promega, E4550) and normalized by total proteins measured with BCA assay (Interchim, UP95424A, UP95425A).

### Electron microscopy

#### Adult Fus^ΔNLS/+^ mice

Mice were anesthetized with intraperitoneal injection of 100 mg/kg ketamine chlorhydrate and 5mg/kg xylazine and transcardially perfused with glutaraldehyde (2.5% in 0.1M cacodylate buffer at pH 7.4) Gastrocnemius muscles were dissected and immersed in the same fixative overnight. After 3 rinses in Cacodylate buffer (EMS, 11650), muscles were post fixed in 0.5% osmium and 0.8% potassium ferrocyanide in Cacodylate buffer 1h at room temperature. Finally, muscles were dehydrated in graded ethanol series and embedded in Embed 812 (EMS, 13940). The ultrathin sections (50 nm) were cut with an ultramicrotome (Leica, EM UC7), counterstained with uranyl acetate (1% (w/v) in 50% ethanol) and observed with a Hitachi 7500 transmission electron microscope (Hitachi High Technologies Corporation, Tokyo, Japan) equipped with an AMT Hamamatsu digital camera (Hamamatsu Photonics, Hamamatsu City, Japan).

#### Embryonic MyoD^iCre/+^; Fus^ΔNLS/ΔNLS^ mice

Dissected Gastrocnemius samples were fixed by immersion in 2.5% glutaraldehyde in Sorenson buffer (0.1M, pH 7.4) for 24 h. The samples were washed and stored in Sorenson buffer. Samples were post fixed in 1% osmium tetroxide in 0.1M cacodylate buffer for 1 hour at 4°C and dehydrated through graded alcohol (50, 70, 90, 100%) and propylene oxide for 30 minutes each. Samples were embedded in Epon 812. Semi-thin sections were cut at 2µm with an ultramicrotome (Leica Ultracut UCT) and stained with toluidine blue, and histologically analyzed by light microscopy. Ultrathin sections were cut at 70nm and contrasted with uranyl acetate and lead citrate and examined at 70kv with a Morgagni 268D electron microscope. Images were captured digitally by a Mega View III camera (Soft Imaging System).

#### Embryonic Fus^+/+^, Fus^ΔNLS/ΔNLS^ and Fus^-/-^ mice

Separated leg samples were fixed first in 2 % glutaraldehyde, 2 % formaldehyde in 0.1 M cacodylate buffer, pH 7.2 for at least 2 hours at room temperature, before the muscle part was dissected. Before embedding the tissue was supported by 2 % LMP agarose and postfixed in 1 % osmium tetroxide including 1.5 % potassium cyanoferrate. Samples were subsequently dehydrated stepwise in ethanol with finally 0.5 % uranyl acetate en-bloc staining during 70 % ethanol. Then the samples were infiltrated in epon. Ultrathin sections of the polymerized sample blocks were cut in different orientations. Representative images were obtained with an electron microscope (Tecnai 12-biotwin, Thermofisher scientific, the Netherlands) with a ditabis plate camera (Ditabis, Pforzheim, Germany).

### Molecular biology and bioinformatics

#### Chromatin immunoprecipitation

Chromatin immunoprecipitation was performed in C2C12 as previously described ^29^. C2C12 were cultured in 6-well plates until 70% of confluency, approximately 1.08 x 107cells were used per conditions. After 48h, cells were fixed with medium containing 1% formaldehyde for 10 min at room temperature. Cross linking was stopped by adding cold glycine at 1.25M during 5 minutes at 4°C. Following washing with PBS1x, cells were lysed with Nuclei lysis buffer (50mM TrisHCL pH8, 2mM EDTA pH8, 0.1% NP40, 10% Glycerol) during 5 minutes then nuclei were sonicated in sonication Buffer (50mM TrisHCL pH8, 10mM EDTA, 0.3% SDS, 1M Na-Butyrate) with Bioruptor (Diagenode, B0102001, 3 Cycles of 5 min - 30s on/ 30s off) to reduce DNA length to between 100 and 500 base pairs ideally 250 base pairs.

Before immunoprecipitation (IP), an input DNA fraction was kept and the protein A/G sepharose beads (Thermo Scientific, 3133) were blocked with10mg/ml tRNA (Sigma, R5636) and 20mg/ml bovine albumin serum (Sigma, A3294) during 2h at 4°C in order to remove unspecific binding. 30µg of sonicated cross-linked chromatin was diluted (3x) with low salt Buffer (50mM TrisHCL pH8, 150mM NaCl, 1mM EDTA, 1% Triton 100X, 0,1% Deoxycholic Acid, 0.1 % SDS, protease inhibitors 1X, 5mM Na-Butyrate), pre cleared during 1h at 4°C with blocked sepharose beads then incubated overnight at 4°C with 5µg of primary antibodies (Fus 294: Bethyl A300-294A, Fus 293: Bethyl A300, A300 293A, RNA pol II: millipore 05-623), or without antibody as negative control. Beads were then resuspended in elution buffer (10 mM Tris pH 8, 1 mM EDTA pH 8), and chromatin was reverse cross linked (0.2 M NaCl, 50 µg/ml of RNase, proteinase K 0.15%) during 3h at 65°C. DNA purification was performed with QIAquick ® PCR Purification Kit (28106 – Qiagen).

#### ChIP-Seq analysis

Reads were aligned to the mouse mm10 genome using bowtie2 ^71^. BigWig coverage tracks were obtained using deepTools bamCoverage. Peak detection was carried out using macs2^72^. Heatmap images were obtained using deepTools computeMatrix and plotHeatmap ^73^. Motif discovery and identification was performed using MEME ^74^ and AME ^75^. To compare coverage and peak files with human datasets, conversion to the hg38 genome was performed using UCSC Genome Browser liftover ^76^. Venn diagram overlaps were generated with ChIPpeakAnno ^77^.

#### Public datasets

Murine ETV5 ^78^, MEF2A ^79,80^ and TAF15 ^81^ Chip-seq datasets were obtained from CistromeDB^82^. Human ETV5, MEF2A and TAF15 Chip-seq binding sites were obtained from ENCODE ^83^. ChIP-Seq signal was retrieved from processed bigwig files and plotted on FUS, ETV5, MEF2a peaks using deepTools computeMatrix and plotHeatmaps

#### RNA-Seq

RNA either from muscle of E18 homozygous *Fus*^ΔNLS/ΔNLS^, *Fus*^-/-^ and control littermates, or from C2C12 cells treated with siRNA against Fus or control siRNA, was extracted using Trizol (Invitrogen). Quality of extracted RNA was determined using the Agilent Bioanalyzer system according to the manufacturer’s recommendations. cDNA libraries after polyA selection were sequenced using an Illumina sequencer with 3-5 biological replicates per condition using single-end 50 bp reads for the mouse muscles samples and 2×150 bp for the C2C12 samples. Reads were aligned to the mouse mm10 genome using STAR ^84^. Differential expression analysis was carried out using edgeR ^85^. Heatmaps were generated in R using pheatmap. Integration of C2C12 and E18.5 muscle datasets was performed using batch correction via limma::removeBatchEffect ^86^, treating each dataset as a batch and using Fus knock-down, Fus knock-in and Fus knock out as bridging populations on the one hand, and WT datasets on the other hand. To integrate ChIP-Seq data, GSEA analyses ^87^ were performed using the gene names of the top 500 ChIP-Seq peaks as genesets, using vsd transformed and batch-corrected expression data.

### Protein biochemistry and phase separation assays

#### Recombinant proteins

FUS recombinant protein was obtained from Novus biologicals (NBP1-42462, immunoprecipitation) or Origene (# TP301808, EMSAs)). ETV1 (# TP310533), ETV4 (# TP760035), ETV5 (# TP761784) and MEF2A (# TP312830) were purchased from Origen. MEF2A-mCherry and ETV5-CFP were produced with the baculovirus expression vector system and purified by the baculovirus platform of IGBMC (Illkirch-Graffenstaden). Expression of recombinant MBP-*tev*-FUS-EGFP-*tev*-His_6_ and His_6_-Tev was performed as described ^42^. In brief, for expression of MBP-*tev*-FUS-EGFP-*tev*-His_6_, the bacterial expression plasmid pMal-*tev*-FUS-EGFP-*tev*-His_6_ was transformed into E.coli BL21-DE3 Rosetta LysS and cells were grown in standard lysogeny broth (LB) medium. At OD600 of ∼0.7-0.8, cells were induced with 0.1 mM IPTG for ca 20h at 12°C. Cells were lysed in resuspension buffer (50 mM Na_2_HPO_4_/NaH_2_PO_4_, pH 8.0, 300 mM NaCl, 10 µM ZnCl_2_, 40 mM imidazole, + 10% glycerol supplemented with 4 mM ≥-mercarptoethanol and 1 µg/ml each of aprotinin, leupeptin hemisulfate and pepstatin A) using sonication and purified by tandem-affinity purification using Ni-NTA agarose (Qiagen) and amylose resin (NEB). The protein was eluted using 300 mM imidazole and 10 mM maltose, respectively and eventually dialyzed into storage buffer (20 mM Na_2_HPO_4_/NaH_2_PO_4_, pH 8.0, 150 mM NaCl, 5% glycerol, 1 mM EDTA, 2 mM DTT).

Expression of His_6_-Tev was induced in E coli BL21 Star using 0.5 −1 mM IPTG at 20°C overnight. Cells were lysed in lysis buffer (50 mM Tris pH 8, 200 mM NaCl, 20 mM imidazole, 10% glycerol supplemented with 4 mM ≥-mercarptoethanol) by addition of lysozyme and sonication in presence of 0.1 mg/ml RNase A. His_6_-Tev was purified using Ni-NTA agarose, washed using lysis buffer containing 1 M NaCl and eluted in lysis buffer containing 600 mM imidazole. The protein was dialyzed against storage buffer (50 mM Tris pH 8, 150 mM NaCl, 20% glycerol, 2 mM DTT). His_6_-3C protease was purified by the IMB protein production core facility and stored in 20 mM Tris pH 7.4, 150 mM NaCl, 20% glycerol, 1 mM DTT.

Protein concentrations for MBP-tev-FUS-EGFP-tev-His_6_ and His_6_-Tev were determined from their absorbance at 280 nm using ε predicted by the ProtParam tool. 260/280 nm ratios of purified proteins were ≤ 0.8.

#### Immunoprecipitation

FUS recombinant protein (1 µg; Novus biologicals NBP1-42462) was incubated overnight at 4°C on a rocking platform in presence of ETV5-CFP (1 µg), MEF2A-mCherry (1 µg) and 20 µl of FUS antibody (Santa Cruz sc-47711) or control IgG antibody (Jackson # 115-007-003) in buffer containing 75 mM Hepes pH 7,3, 100 mM KOAc, 1% NP-40, 2,5% glycerol, 0,5 mM DTT and PIC 1X. A/G magnetic bead (20 µl - ThermoFisher Scientific #88802) was added to the sample and placed on a rocking platform at room temperature during 1h30 then the supernatant was removed and the beads were washed three times with RIPA. Finally, beads were resuspended in Laemmli sample buffer and protein were eluted by boiling 10 minutes at 90°C and analyzed by SDS-PAGE and immunoblotting using FUS (Santa Cruz sc-47711), ETV5 (Proteintech 13011-1-AP) and MEF2A (Santa Cruz sc-17785) antibody.

#### EMSA

Oligonucleotides were purchased from Eurogentec. For MRPL30 probe, a single strand sense (FAM-labelled; 5’-FAM-TTTTAACGGCGTGGTTTGAATGTTGCACTGGGTCCACCTGCCTCTGTCTCCGGCTGCGGGGACGTTCGGA-3’) and antisense (5’-TCCGAACGTCCCCGCAGCCGGAGACAGAGGCAGGTGGACCCAGTGCAACATTCAAACCACGCCGTTAAAA-3’) oligonucleotides were annealed in buffer containing 100 mM Tris, pH 7.5-8, 500 mM NaCl, 10 mM EDTA at 90°C for 5 minutes and gradually cooled at room temperature during 45 min. MRPL30 mutant probe was obtained by annealing single strand sense (FAM-labelled; 5’-FAM-TCCGAACGTCCCCGCAGCCGGAGACAGAGGCAGGTGGACCCAGTGCAACATTCAAACCACGCCGTTAAAA-3’) and antisense (5’-TTTTAACGGCGTGGTTTGAATGTTGCACTGGGTCCACCTGCCTCTGTCTCCGGCTGCGGGGACGTTCGGA-3’) oligonucleotides.

EMSAs were performed in 20 µl reaction volume containing 20 mM HEPES-KOH pH 7.5, 5 mM MgCl2, 50 mM KCl, 10 mM DTT, 10% Glycerol. Binding reactions were performed in presence of increasing concentration ETV5 (60, 120, 180 and 240 nM) or constant concentration of ETV1, ETV4 or ETV5 (120 nM) and/or Fus (300 nM) and/or MEF2A (50 nM) and MRPL30 double strand probe WT or mutant (5 nM/reaction). After 25 min incubation at 25°C, the reaction mixture was loaded onto a 6% polyacrylamide gel (ThermoFisher Scientific # EC63652BOX) in 0.5x TBE and electrophoresed at 100 V for 2–3 h. The gels were imaged by iBright FL1000 Invitrogen Imaging System and the signal intensity was quantified using the iBright™ Analysis Software.

#### Microscopy condensation assay

Purified MBP-*tev*-FUS-EGFP-*tev*-His_6_, MBP-3C-CFP-ETV5 and MBP-3C-MEF2A-mCherry were mixed at concentrations determined ideal for DNA binding by EMSA in 50 mM Tris pH 7.5 supplemented with 2 mM DTT containing 150 mM NaCl final. Cleavage of the MBP-solubility tag was induced by addition of 1.3 µM His_6_-Tev and/or 1.6 µM His_6_-3C proteases, respectively, in 18-well ibiTreat dishes (Ibidi) and imaged by confocal microscopy using a Leica Stellaris Microscope (Imaging Core Facilty of IMB Mainz) after 45-60 min. Images were acquired using unidirectional scanning at a pixel size of 64 nm using a HC PL APO CS2 63x/1.4 oil objective. Using a white light laser, CFP-ETV5 was excited at 440 nm and detected using a HyDX2 detector (445-742 nm), FUS-EGFP was excited at 489 nm and detected by a HyDS3 detector (494-593 nm), MEF2A-mCherry was excited at 587 nm and detected by a HyDX4 detector (593-750 nm). Images were analyzed using Fiji/Image J built in plug-ins for particle analysis. In brief, first a median filter (3 px) and subsequently a common threshold to include all condensates were applied to all images from the same experiment. Finally, images were converted into binary and watershedding was performed if applicable. Eventually, condensates were analyzed using the “Analyze Particles” plugin.

### Statistics

All results are presented as mean ± standard error of the mean (SEM) and differences were considered significant when p < 0.05. Significance is presented as follows: * p<0.05, ** p<0.01, and *** p<0.001.

For comparison of two groups, two-tailed unpaired Student’s *t* –test was used in combination with F-test to confirm that the variances between groups were not significantly different.

Comparison for more than two groups was performed using one-way ANOVA and Tukey or Bonferroni’s multiple comparison *post hoc* test or Kruskal Wallis as indicated in figure legends. For cell biology and biochemistry experiments, all experiments were performed at least 3 times, each biological replicates consisting in several technical replicates. In these cases, we used a Nested one-way ANOVA with Tukey’s multiple comparisons test, pairing technical replicates of each biological replicate experiment.

For statistical analysis of dorsal abdominal muscle number in *Drosophila* pharate adult flies, two-way ANOVA with Tukey’s multiple comparisons test and Kruskal-Wallis test with Dunn’s multiple comparisons test were used. To analyze muscle misorientation, Wilcoxon signed rank test was used to evaluate differences as compared to the control genotype, and direct comparison between MHC-GAL4, *caz^FRT^* and MHC-GAL4, *caz^FRT^*; UAS-FLP was analyzed by Mann-Whitney test. Chi-square statistics was used to analyze offspring frequency data. For life span analysis, the log-rank test was used to test for statistical significance. Motor performance was analyzed using the Mann-Whitney U test to compare climbing speed of individual flies per genotype and per run. Because all flies were tested in three independent runs, three P values were generated per genotype. These P values were combined using the Fisher’s combined probability test. Statistical analysis was performed using Graphpad Prism software.

## Acknowledgments

We thank Marc David Ruepp for advice with LLPS deficient FUS mutants, Peter C Hollenhorst for helpful advice on interactions between FET proteins and ETS-transcription factors, Todd Gulick and Ron Prywes for providing expression and reporter plasmids, and Karina Mildner, Claudia de Tapia, Marion Boutry and Aurélie Bombardier for technical assistance. This work was funded by Agence Nationale de la Recherche (ANR-16-CE92-0031, ANR-19-CE17-0016, ANR-20-CE17-0008, ANR-21-CE17-0039), by the Interdisciplinary Thematic Institute NeuroStra, as part of the ITI 2021-2028 (Idex Unistra ANR-10-IDEX-0002, ANR-20-SFRI-0012), by Fondation Bettencourt (Coup d’élan 2019 to LD), Fondation pour la recherche médicale (FRM, DEQ20180339179), Axa Research Funds (rare diseases award 2019, to LD), Association Francaise de Recherche sur la sclérose latérale amyotrophique (2016, 2021 to LD and ES), Radala Foundation for ALS Research (to LD and ES), the Association Française contre les Myopathies (AFM-Téléthon, #23646 to LD and ES), TargetALS (to LD) JPND (HiCALS project, to LD), ‘Stichting ALS Nederland’ (to ES), the ‘Prinses Beatrix Spierfonds’ (W.OR22-03, to ES), the Muscular Dystrophy Association (MDA 946876, to ES) and an NWO Open Competition ENW-M grant (to ES) and the German Research Foundation (DFG) within the Heisenberg Programme (project number 442698351, to DD). E.S. is supported by an ERC consolidator grant (ERC-2017-COG 770244). LD is USIAS fellow 2019. We are grateful to IGBMC Baculovirus platform, PICSTRA Imaging platform (UMS38, CRBS), PEFRE animal facility platform (UMS38, CRBS), IMB Protein Production and Microscopy Core Facilities for technical assistance and support; Stellaris Confocal Microscope at IMB Microscopy Core Facility is funded by the Deutsche Forschungsgemeinschaft (DFG; German Research Foundation project number 497669232).

