## Supplementary Figures for "FUS controls muscle differentiation and structure through LLPS mediated recruitment of MEF2 and ETV5"

Picchiarelli, Wienand et al.

### Supplementary Figures

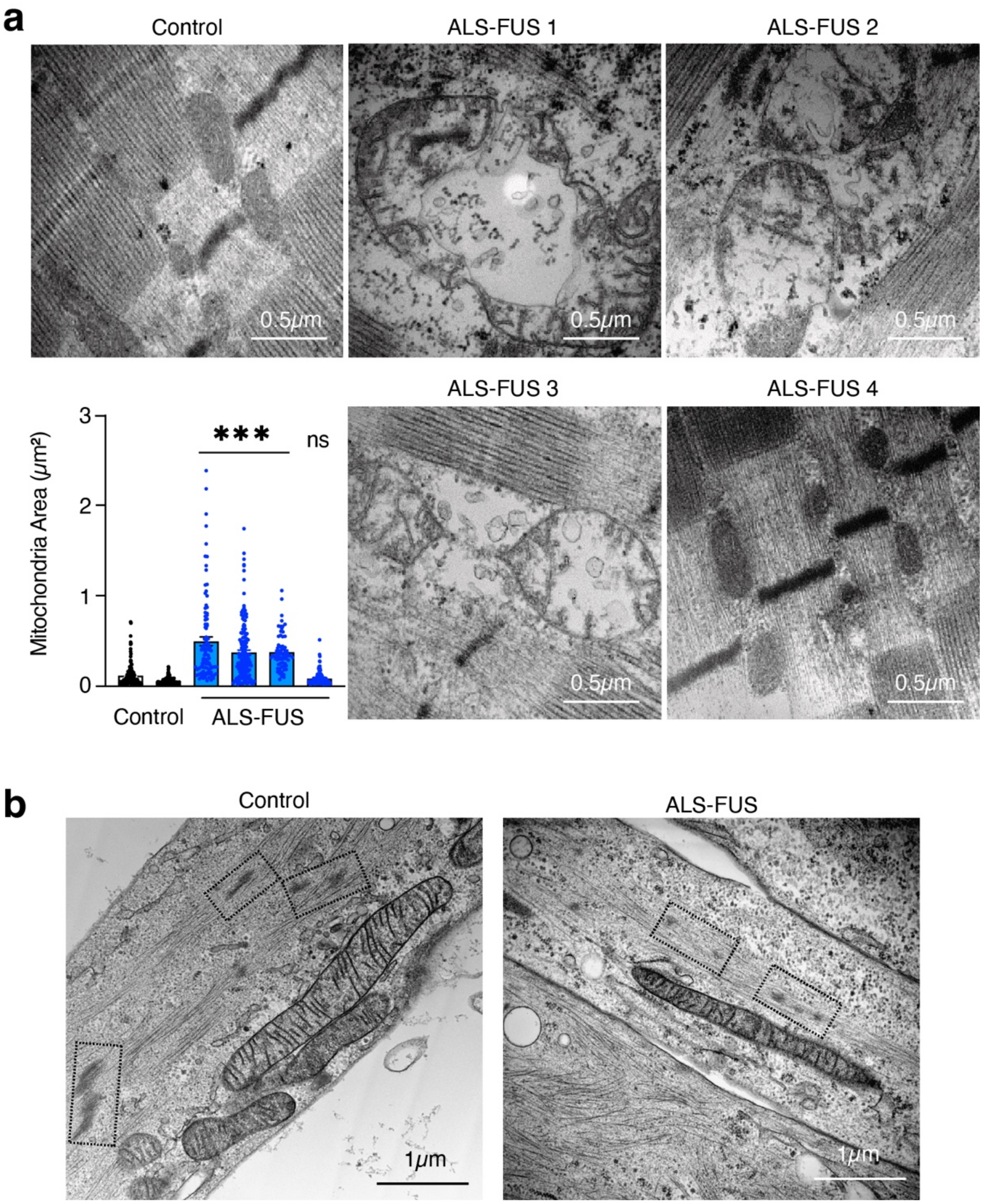


**Supplementary Figure 1: Abnormal ultrastructure in *FUS*-ALS muscles.**

A: Electron micrographs of muscle mitochondria in healthy control showing normal size and structure of mitochondria, and in *FUS*-ALS patients, showing fragmentation, swelling or complete destruction of mitochondria in FUS1, FUS2, and FUS3 patients, while FUS4 patient did not show altered mitochondrial ultrastructure.

Lower left panel shows quantification of mitochondrial surface area with a significant increase in mitochondrial size in *FUS*-ALS patients, except for patient FUS4 (ns: not significant). One-way ANOVA with Tukey's multiple comparisons test: Control versus FUS1, FUS2, FUS3: ***, p<0.0001. Scale bars: 0.5µm.

b: Ultrastructural alterations in iPSC-derived myotubes**.** Electron micrographs of sphere-derived myotubes from isogenic control and *FUS*-ALS patient iPSC lines, showing the mitochondria and Z-lines (dashed boxes). Z-lines in mutant myotubes appear less dense than in the control.

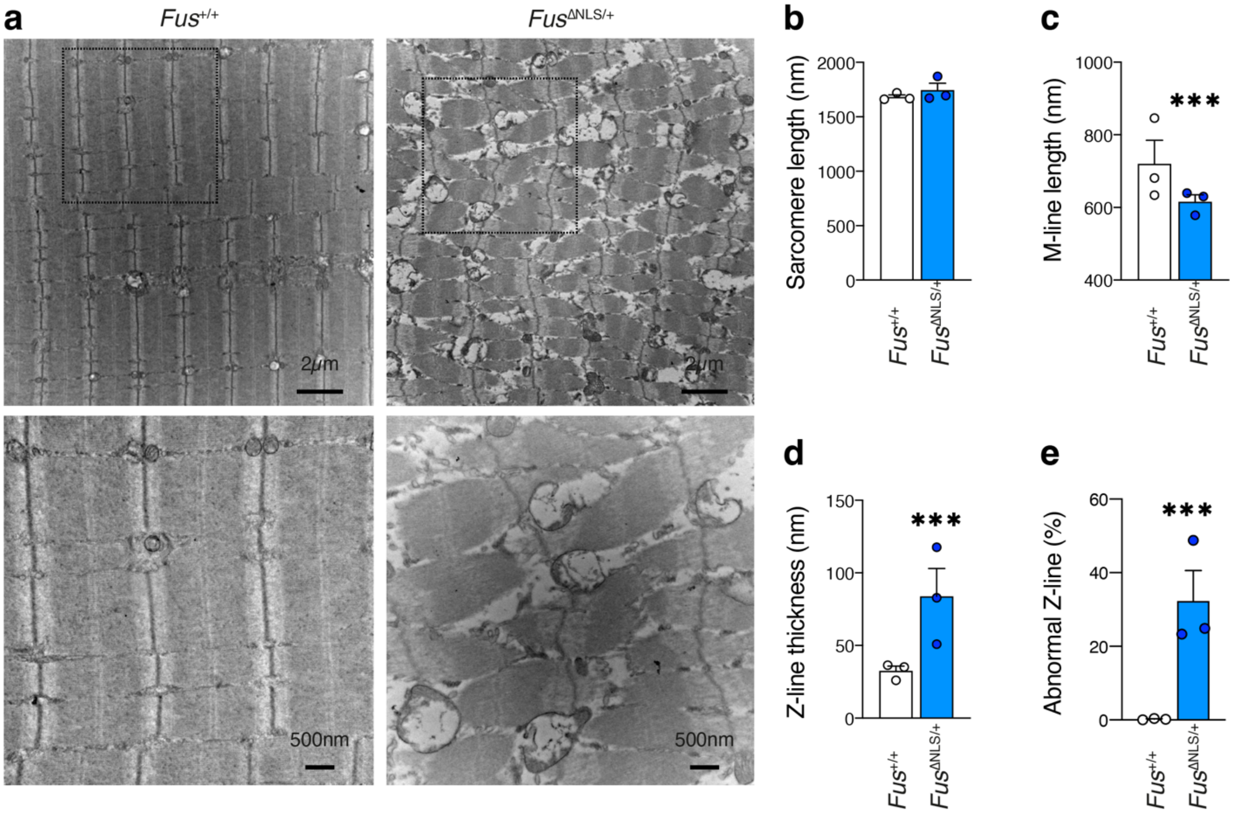


**Supplementary Figure 2: Heterozygous *Fus* mutation in mouse leads to muscle ultrastructural defects**

a: Representative TEM images in gastrocnemius muscle of 22-month-old *Fus*$^{\Delta NLS/+}$ mice and their control littermates. A higher magnification of the indicated region (dashed box) is shown in the lower panels.

b-e: Quantitative analyses of ultrastructural parameters of sarcomeres in adult *Fus*$^{\Delta NLS/+}$ mice and their control littermates. Dots represent individual animals analyzed (n=3), 20 pictures were analyzed per animal. ***P<0.001 by nested t-test.

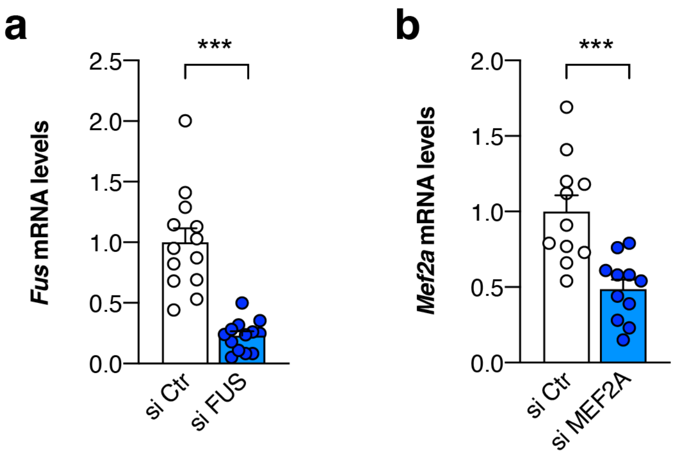


**Supplementary Figure 3: qPCR validation of siRNA knockdown**
a-b: C2C12 cells were transfected with si Ctr, si FUS (A) or si MEF2A (B) and RNA was extracted 24h after transfection. *Fus* (a) or *Mef2a* (b) mRNA levels were measured using RT-qPCR. ***P<0.001 by Student's t-test.

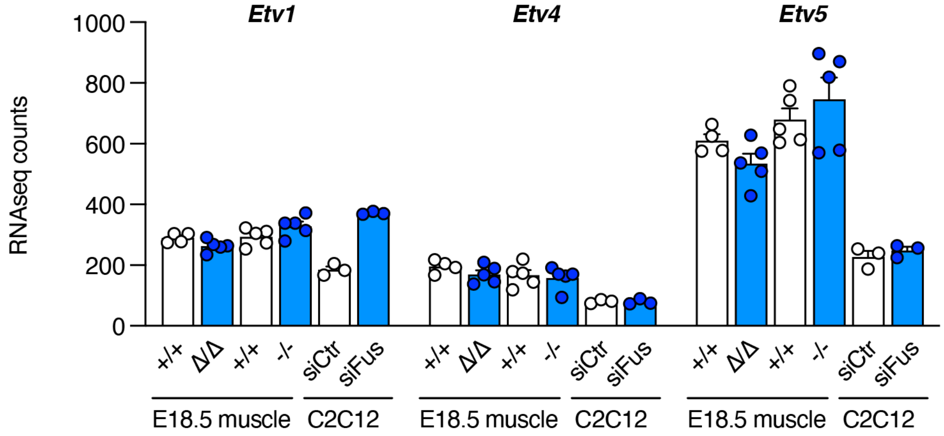


**Supplementary Figure 4: normalized RNAseq counts for Etv1, 4 and 5 in various models**

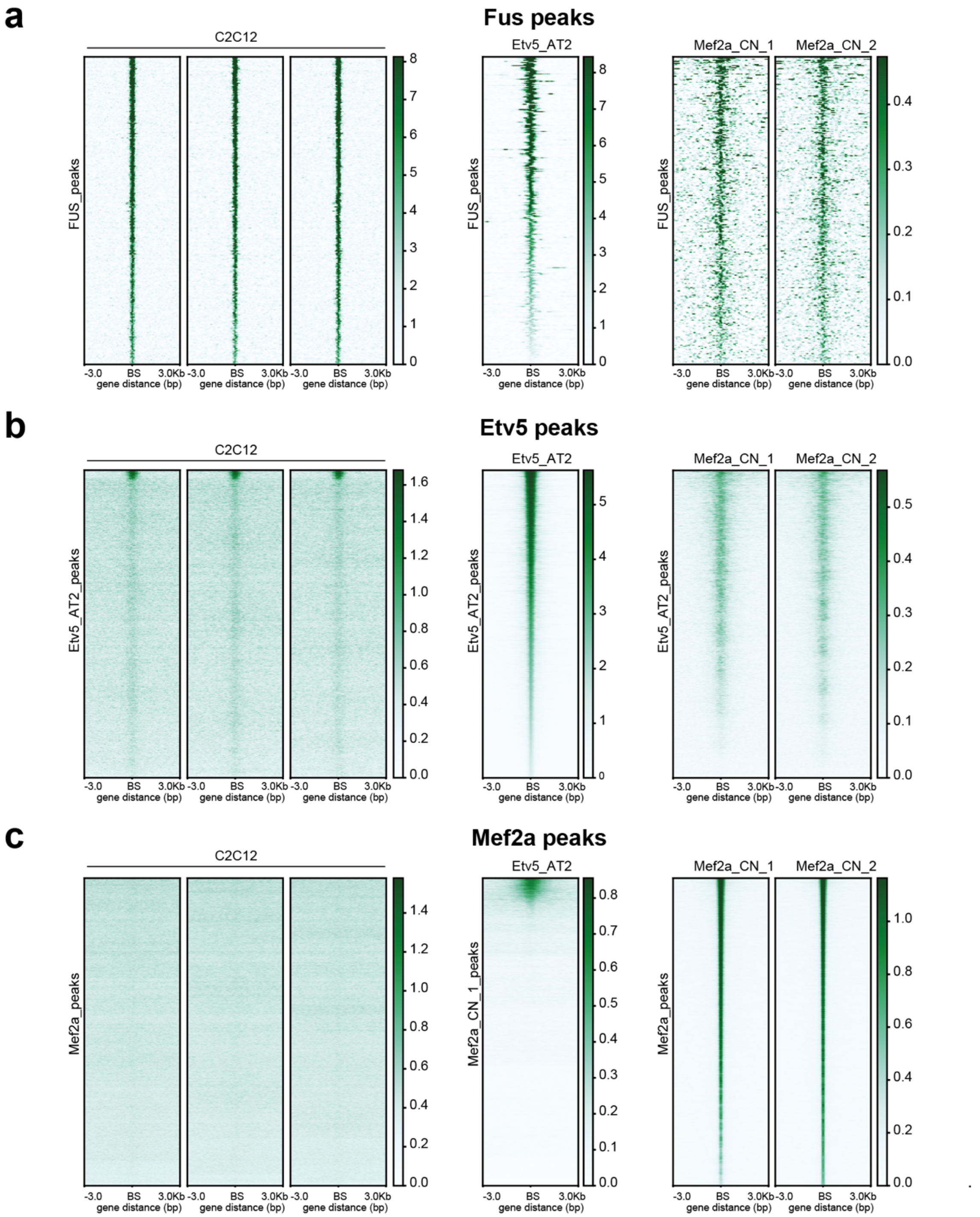


**Supplementary Figure 5: Colocalization of FUS, ETV5 and MEF2A on FUS peaks.**
Heatmaps showing FUS (a), ETV5 (b) and MEF2A (c) peaks in the 3 replicates of C2C12 ChIP seq. Note colocalization of all three factors on FUS peaks only. ETV5 and MEF2A peaks were predicted using publicly available datasets in mouse AT2 cells and mouse cortical neurons (CN) respectively. References of the datasets are indicated in methods.

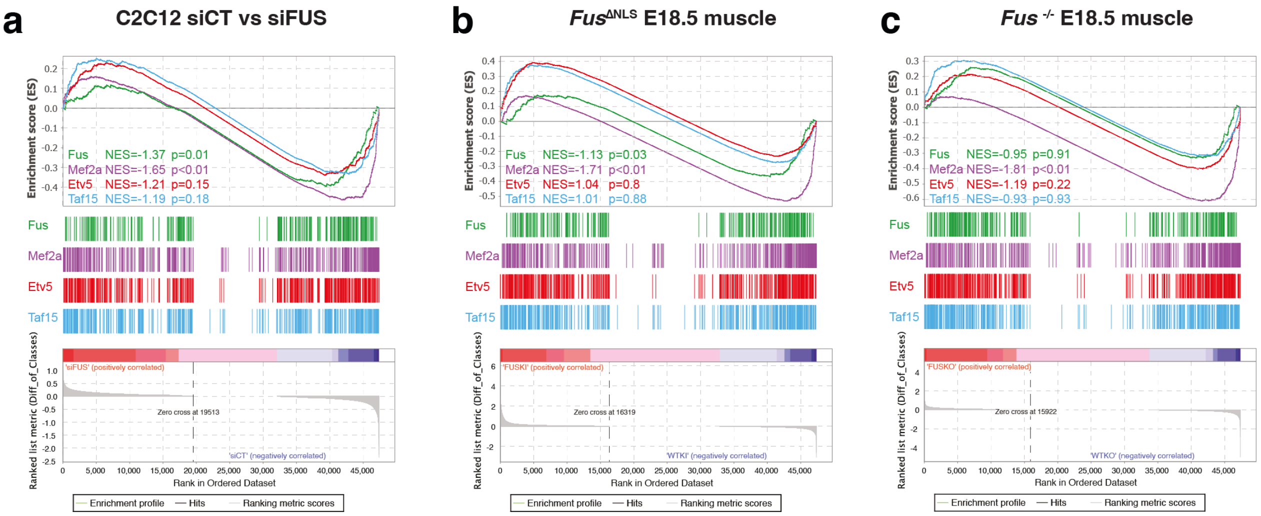


**Supplementary Figure 6: Correlation between FUS, co-factor binding and gene expression fold change.**

Gene set enrichment analysis (GSEA) using genes closest to FUS (green), MEF2A (purple), ETV5 (red) and TAF15 (teal) peaks using gene expression data from C2C12 siFus vs siCT (a), E18.5 muscle *Fus*$^{\Delta NLS/\Delta NLS}$ vs *Fus*$^{+/+}$ (b), E18.5 muscle *Fus*$^{-/-}$ vs *Fus*$^{+/+}$ cells (c).
Note significant correlations for FUS in siFus vs siCT in C2C12 cells and E18.5 muscle from *Fus*$^{\Delta NLS/\Delta NLS}$ vs *Fus*$^{+/+}$ and for MEF2A for all three datasets.

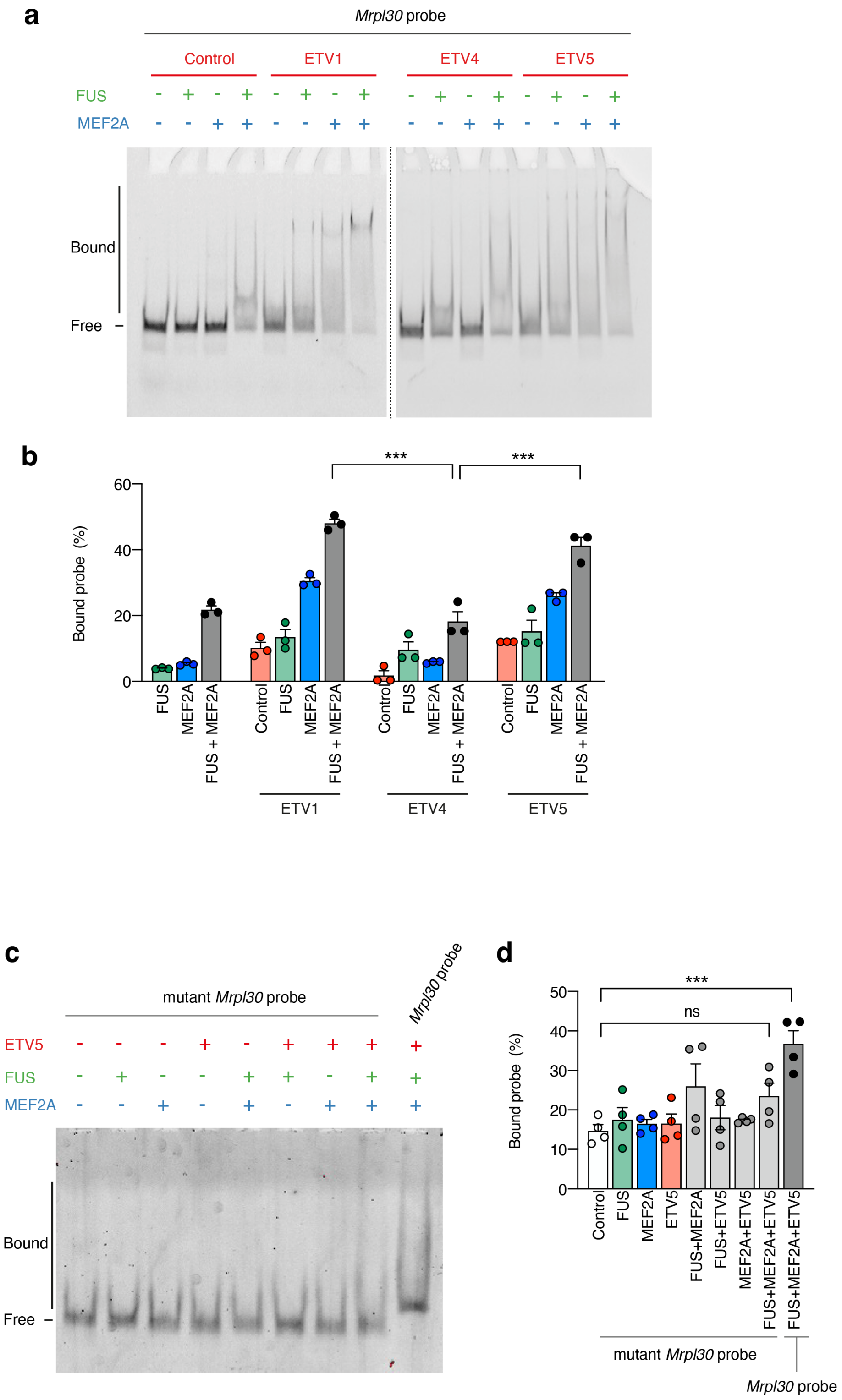


**Supplementary Figure 7: Additional EMSA specificity controls**

a: Cooperative binding of FUS, MEF2A and the different ETV factors (ETV1, ETV4 or ETV5) to FAM labelled *Mrpl30* promoter-derived DNA probe. EMSA experiments were performed by using a constant concentration of FUS (300 nM), MEF2A (50 nM) and ETV (120 nM) and 5 nM of MRPL30 DNA probe.

b: Graph showing the percentage of *Mrpl30* probe bound. The intensity of the signals of the protein-DNA complex were quantified using the iBright™ Analysis Software. Error bars represent SEM of three independent experiments.
c: EMSA experiments were realized with a constant concentration of FUS (300 nM), MEF2A (50 nM) and ETV5 (120 nM) and 5 nM of *Mrpl30 mutated* promoter-derived DNA probe where the ETV5 binding motifs have been mutated. As positive control, the last lane was performed with the *Mrpl30* promoter-derived DNA probe not mutated.
d: Graph showing the percentage of *Mrpl30* mutated or control probe bound. The intensity of the signals of the protein-DNA complex were quantified using the iBright™ Analysis Software. Error bars represent SEM of four independent experiments.

**a** **Prediction of naturally disordered regions**

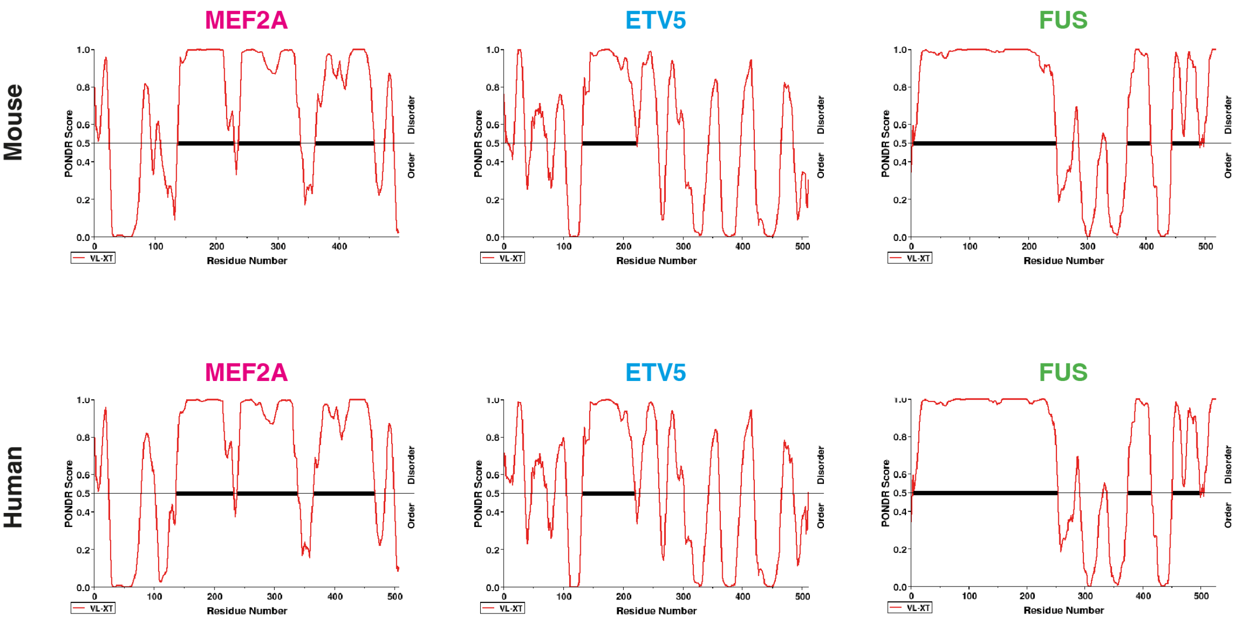


**b** **Prediction of phase separation behavior**

| Protein | PSP Score | Phase separation prediction |
|---|---|---|
| Human FUS | 0.9987 | Yes |
| Mouse FUS | 0.9987 | Yes |
| Human MEF2A | 0.969 | Yes |
| Mouse MEF2A | 0.9399 | Yes |
| Human ETV5 | 0.8531 | Yes |
| Mouse ETV5 | 0.7067 | Yes |

**Supplementary Figure 8: FUS, MEF2A and ETV5 LLPS predictions**
a: Results of PONDR prediction for mouse and human MEF2A, ETV5 and FUS proteins. All 6 proteins display intrinsically disordered regions.
b: Prediction of phase separation behavior using PSP. All 6 proteins are predicted to undergo phase separation.

**Supplementary Tables**

**Supplementary Table 1: muscle biopsies from ALS patients and controls**

| **Name** | **Gender, Age*** | ***FUS* mutation** | **Electromyography** |
|---|---|---|---|
| CNTL1 | Female, (33) | n.a | |
| CNTL2 | Male, (30) | n.a | |
| FUS1 | Male, 26 (26) | R495QfsX527 | Intensive PSA, neurogenic and myogenic MUAP |
| FUS2 | Female, 24 (27) | K510R | Intensive PSA, neurogenic MUAP |
| FUS3 | Male, 39 (42) | K510R | Moderate PSA, myogenic and far less neurogenic MUAP |
| FUS4 | Male, 24 (24) | G478L | Moderate PSA, neurogenic and myogenic MUAP |

*Age of onset (age at biopsy); CNTL: Control, PSA: pathological spontaneous activity; MUAP: motor unit action potential
